# Directional reorientation of migrating neutrophils is limited by suppression of receptor input signaling at the cell rear through myosin II activity

**DOI:** 10.1101/2021.04.04.438336

**Authors:** Amalia Hadjitheodorou, George R. R. Bell, Felix Ellett, Shashank Shastry, Daniel Irimia, Sean R. Collins, Julie A. Theriot

## Abstract

To migrate efficiently to target locations, cells must integrate receptor inputs while maintaining polarity: a distinct front that leads and a rear that follows. Here we investigate what is necessary to overwrite pre-existing front/rear polarity in neutrophil-like HL60 cells migrating inside straight microfluidic channels. Using subcellular optogenetic receptor activation, we show that receptor inputs can reorient weakly polarized cells, but the rear of strongly polarized cells is refractory to new inputs. Transient stimulation reveals a multi-step repolarization process, confirming that cell rear sensitivity to receptor input is the primary determinant of large-scale directional reversal. We demonstrate that the RhoA/ROCK/myosin II pathway limits the ability of receptor inputs to signal to Cdc42 and reorient migrating neutrophils. We discover that by tuning the phosphorylation of myosin regulatory light chain we can modulate the activity and localization of myosin II and thus the amenability of the cell rear to ‘listen’ to receptor inputs and respond to directional reprogramming.

## INTRODUCTION

Neutrophils, the most abundant circulating leukocytes in humans, comprise the first line of innate immune defense. Their directed migration is mediated by detection of chemoattractant such as fMLF via G protein-coupled receptors (GPCRs), fairly evenly distributed on the cell surface^1, 2^. Ligand binding to the receptor activates signaling via G_i_, leading to cell polarization and directional motility^3^.

An early manifestation of neutrophil polarization is the generation of steep antagonistic gradients of Cdc42 and RhoA activity, formed synchronously towards the cell front and rear, respectively^4^ (Fig. 1a). At the front, Cdc42 and Rac1 induce actin polymerization, while RhoA regulates myosin II contractility at the rear. The front and rear signaling modules are mutually exclusive^3^ and are governed by positive feedback loops for self-amplification and polarity maintenance^5–7^. In addition, tension by the plasma membrane has been demonstrated to act as a long-range inhibitor, mechanically preventing the generation of multiple cell fronts^8^.

**Figure 1:**
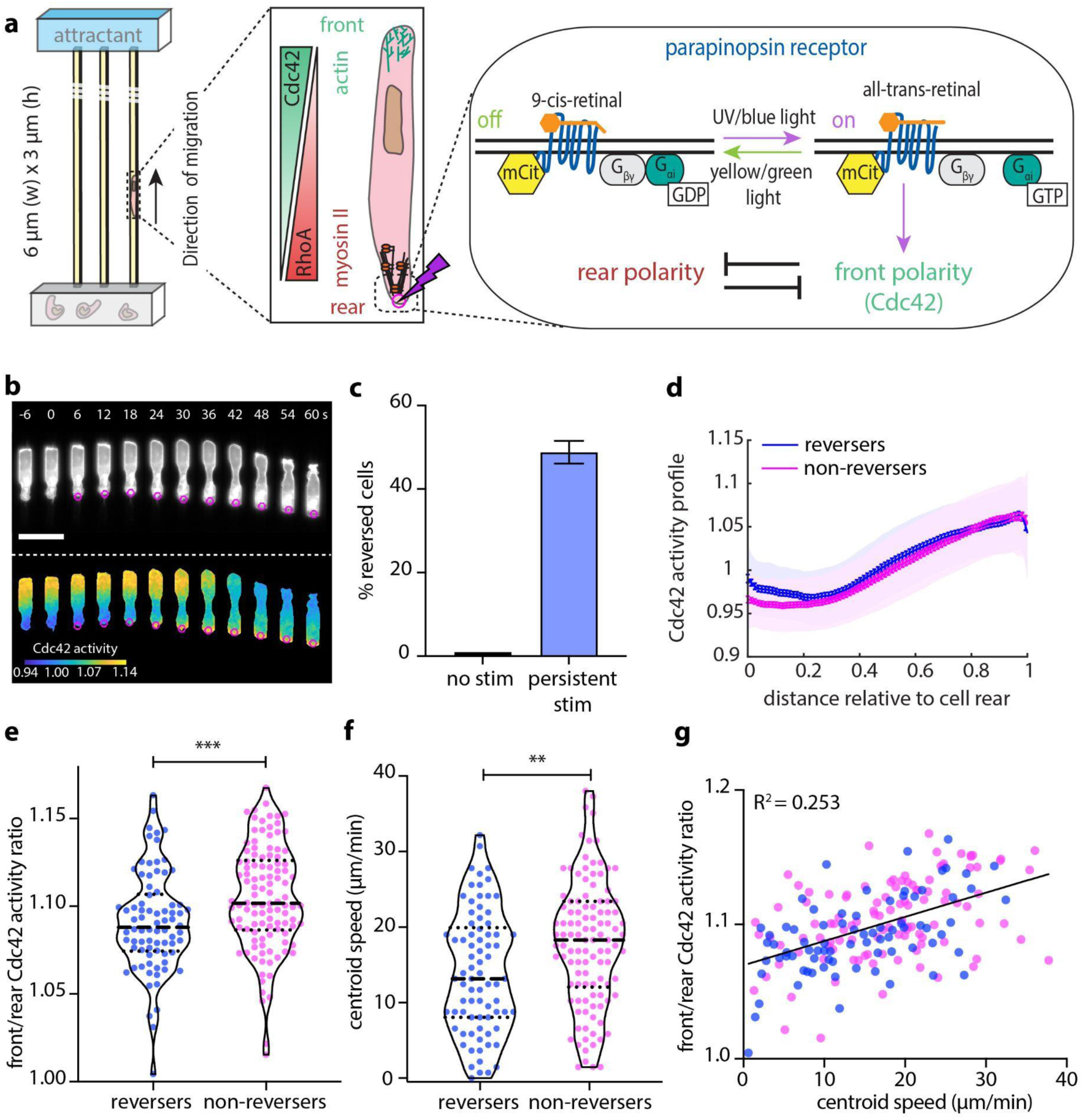
Persistent optogenetic stimulation is sufficient to reverse weakly polarized and slowly migrating cells. **(a)** Schematic representation of the front-rear polarity reversal assay. HL60 cells expressing parapinopsin, an optogenetic GPCR, migrate inside microfluidic devices that harbor straight channels. A heat inactivated serum gradient is used as an attractant. The opsin can be turned on using UV/blue light and off with yellow/green light, offering complete control of the receptor activity. Blue light stimulation (magenta circle/lighting bolt) at the rear of the cell turns on the receptor. The receptor is coupled to the polarity signal transduction network and activation re-enforces the front module. **(b)** Live-cell imaging snapshots of cells expressing parapinopsin and a red/far-red Cdc42 FRET sensor. Cells migrate unperturbed for 60 s prior to initiation of persistent pulsed stimulation (magenta circles) at their rear. Upper and lower panels show the registered sensors (grey scale) and the computed Cdc42 activity, respectively. Images captured every 3 s and subsampled for illustration purposes. Scale bar: 25 µm. **(c)** Box plot of the percentage of cells that reversed with persistent pulsated stimulation (n=336 cells from 36 independent experiments) versus non-stimulated control (n=32 cells from 3 independent experiments). Error bar represents confidence intervals assuming a binomial distribution around the cumulative mean. **(d)** Mean Cdc42 activity profiles for n=78 reverser (blue) and n=110 non-reverser (magenta) cells averaging cell profiles over 15 s prior to stimulation (lines: means, shaded regions: SD, error bars: SE). **(e-f)** Violin plots of mean front/rear Cdc42 activity ratio **(e)** and of mean centroid speed **(f)** of n=78 reversers and n=110 non-reversers cells, averaging over 15 s prior to stimulation; p-values of two-sided Wilcoxon rank sum test (**: p<0.01, ***: p<0.001). **(g)** Linear regression between calculated mean front/rear Cdc42 activity, shown in **(e)**, and mean centroid speed, shown in **(f)**, prior to stimulation, parametric Pearson correlation yields R^2^=0.253.

Once established, front-rear polarity in migrating neutrophils is thought to be relatively stable, as evidenced by the observation that polarized neutrophils often steer their original front to make a U-turn instead of repolarizing, suggesting that neutrophils are more sensitive to chemoattractants towards their front as compared to their rear^1^. To examine the role of polarity in chemoattractant sensing, researchers have historically challenged neutrophils migrating on 2-D planar substrates using point sources of chemoattractant at different angles with respect to the original direction of migration^3, 9–11^. In these experiments, chemoattractant is typically delivered using a micropipette positioned near the cell, resulting in diffusion of the attractant over the entire cellular surface. Thus, it is difficult to decouple whether the rear is intrinsically less responsive to chemoattractant signaling than the front, or whether the greater amplification of signaling inputs at the cell front gave it the advantage. Recent studies have leveraged microfluidic devices, examining the response of neutrophil-like HL60 cells and *Dictyostelium* cells to flipping the direction of the chemoattractant gradient when cells are confined in 1-D channels and cannot physically perform a U-turn^12, 13^.

To probe the cellular sensitivity to receptor inputs at the level of signaling, and to determine what is necessary to overwrite front-rear polarity, we used an optogenetic approach to locally activate G_αi_ signaling, independent of fMLF, and drive cells to dynamically repolarize without modifying the environment in which they are embedded. We found that persistent optogenetic receptor activation at the rear of neutrophils migrating in 1-D microfluidic channels is sufficient to reorient weakly polarized and slowly migrating cells. However, in more strongly polarized cells, myosin II and RhoA activity limit the ability of the rear to respond, even at the level of signal transmission to Cdc42, creating a cell rear that is refractory to new receptor inputs. We show that by tuning the phosphorylation of myosin regulatory light chain, we can modulate the activity and localization of myosin II and thus the amenability of the cell rear and the ability of cells to reverse their direction of motion.

## RESULTS

### Persistent optogenetic stimulation is sufficient to reverse weakly polarized and slowly migrating cells

We generated a neutrophil-like HL60 cell line stably expressing parapinopsin (a light-sensitive G_αi_-family GPCR) and a tdTomato/tdKatushka2 Cdc42 FRET biosensor, spectrally compatible with the reversible parapinopsin stimulation. This system allows direct recording of downstream Cdc42 activity in cells whose migration is guided by parapinopsin^14^. We confined cells to migrate inside 1-D straight microfluidic channels in response to spatial serum gradients (Fig. 1a). Using a 407 nm laser, we locally stimulated cells with 10 ms light pulses and the spatial precision of about a 1 µm diameter spot, using real-time image analysis to automatically and dynamically position the activation spot at the appropriate location (see Methods). We found that repeatedly delivering light pulses every 3 s at the cell rear was sufficient to overwrite the front/rear polarity, drive a chemotaxis-like response on the level of Cdc42, and ultimately reverse the direction of motion in a subset (about 47%) of cells (Fig. 1b & Movie S1).

We were initially surprised by the observation that just 47% of cells reversed (Fig. 1c). To better understand why only some cells were able to reverse direction, we first used flow cytometry to measure the mCitrine-tagged parapinospin receptor (abbreviated as mCit on Fig. 1a) and found that over 97% of cells expressed the construct (Supplemental Fig. 1a). We next confirmed that at least 70-93% of cells were responsive to a 5-pulse optical stimulation administered at their center (Supplemental Figs. 1b-1c & Movie S2). These estimates of inherently responsive cells were significantly higher than the 47% of reversing cells, suggesting that the reversals were suppressed due to some other kind of variation among cells.

We then examined whether there were any measurable pre-stimulation differences between reversing and non-reversing cells. Reversing cells tended to be weakly polarized (Figs. 1d-1e), and slower migrators (Fig. 1f), although there was considerable overlap between the distributions of reversing and non-reversing cells. The primary determinant of reversibility came from the cell rear (Supplemental Fig. 2a-2c). Mean front/rear Cdc42 activity ratio appeared to correlate with centroid speed (Fig. 1g). We found no correlation between a cell’s position along the channel and its likeliness to repolarize upon stimulation (Supplemental Fig. 2d).

All together our analysis suggested that pre-existing variations in cell state could be influencing the ability of the cell to respond to new receptor inputs, and that the amenability of a cell to reverse polarity may be tunable by the sensitivity of the cell rear.

### In reversing cells Cdc42 activation at the stimulated rear begins immediately and cells reverse their direction of migration before Cdc42 activity flips polarization

To gain further insight into what may be controlling the rear sensitivity to receptor inputs, we studied in a stepwise mechanistic manner the order of events during optogenetic-driven cell reversals. We quantified the Cdc42 activity at the original rear and front for 78 reversing cells (Fig. 2a) as well as the derivative of Cdc42 activity at each edge (Fig. 2b). Notably, Cdc42 activity began to increase at the stimulated rear almost immediately, whereas the decrease in Cdc42 activity at the old front had a much slower onset. Most strikingly, the direction of migration flipped 27 s post-stimulation (Fig. 2c), considerably before the two cell edges reached equal levels of Cdc42 activity (Cdc42 cross-point) 51 s post-stimulation. Thus, the cell edge that has more Cdc42 activity is not necessarily the driving front.

**Figure 2:**
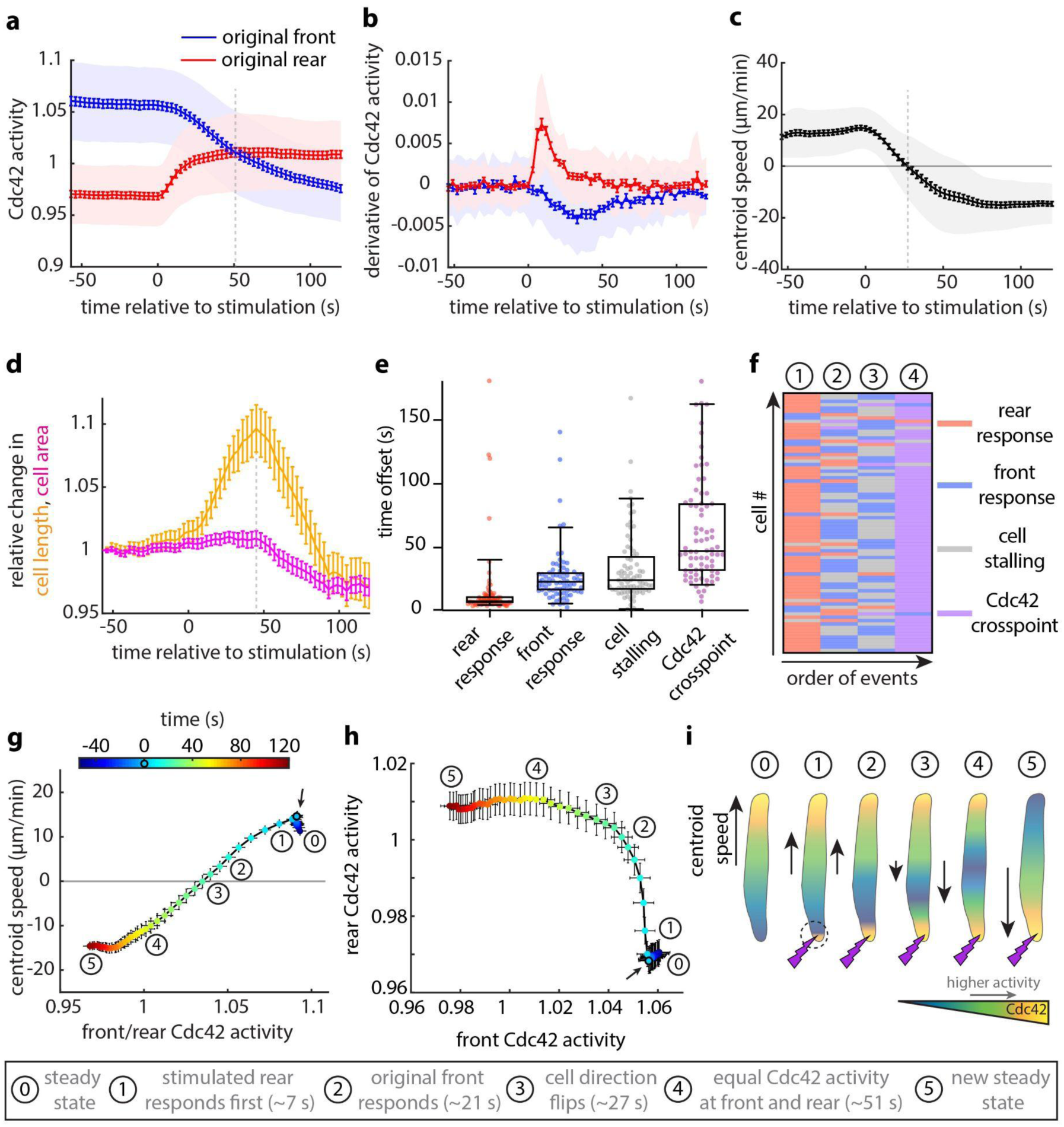
In reversing cells Cdc42 activation at the stimulated original rear begins immediately and cells reverse their direction of migration before Cdc42 activity flips polarization. **(a-b)** Mean magnitude **(a)** and derivative **(b)** of Cdc42 activity at the original cell front (blue) and rear (red) over time (lines: means, shaded regions: SD, error bars: SE). Data are averages from n=78 reverser cells from 36 independent experiments. Dashed vertical line in **(a)** shows Cdc42 crossover point. **(c)** Centroid speed of cells that reversed their migratory direction over time (line: mean, shaded region: SD, error bars: SE). Data are averages from the same 78 cells shown before. Dashed vertical line indicates when cell stalling (zero speed) occurs. **(d)** Relative change in cell length (orange) and cell area (magenta) both normalized to the initial length and area for each of 78 cells. Dashed vertical line indicates the time when the relative change in cell length reaches maximum value. Note that the normalized area appeared to decrease post-stimulation, overshooting below the starting value. This relative decrease in cell area was associated with a smaller rear after repolarization. Moreover, the Cdc42 activity at the new emerging front was restricted in a smaller depth as compared with the pre-stimulus steady state (Movie S1), suggesting that the repolarized cell adopted a sharper Cdc42 activity profile with a more constricted rear. **(e)** Box plots of the time-offset for rear response, front response, cell stalling, and Cdc42 cross-point for 78 cells that reversed. **(f)** Heat map of the order of events (columns) across 78 reversed cells (rows). Circled numbers correspond to the cell states shown in **(i)**. **(g-h)** Phase diagrams of mean centroid speed versus mean front/rear Cdc42 activity **(g)** and of mean rear versus mean front Cdc42 activity **(h)**, averaging over 78 reversed cells. Colored points correspond to the average cell state at different time points relative to the start of the persistent pulsated stimulation (black circle/arrow). Circled numbers indicate the cell states as summarized in **(i)** and error bars denote SE. **(i)** Cartoon depiction of the average order of events for a cell that reverses.

Moreover, as the stimulated original rear began to increase local Cdc42 activity, the two cell edges pulled in competing directions, causing the normalized cell length to increase and peak 45 s after the initiation of stimulation (Fig. 2d), temporally close to the Cdc42 cross-point. The normalized cell area remained constant during this tug-of-war between the original and emerging front (Fig. 2d). We conclude that the stimulated rear quickly assumed front character and started to move towards the opposite direction, before the signal was transmitted to the original front. In others words, flipping polarity was not instantaneous, as flipping a switch, but was a multistep transition.

The above order of events was also evident in single cells (Fig. 2e-2f). Phase diagrams of mean centroid speed versus mean front/rear Cdc42 activity ratio (Fig. 2g) and of mean rear versus mean front Cdc42 activity (Fig. 2h) further illustrated the transitional states a cell occupied before flipping its polarity and resuming a new steady state (Fig. 2i). This analysis confirmed that Cdc42 activity at the stimulated rear rose quickly, and only after the Cdc42 activity at the original rear got quite high did the Cdc42 activity at the original front start decreasing.

### Transient stimulation reveals a variety of distinct cellular responses

Surprised by the observation that cells reversed their direction of migration before Cdc42 activity flipped polarization, we asked what would happen if we stopped the stimulation in between these two key events, namely after the cell stalling at 27 s and before the Cdc42 cross-point at 51 s. We found that transient 12-pulse stimulation resulted in four distinct classes of observed cellular responses that manifested differently on the level of Cdc42 activity with distinct migration patterns (Fig. 3a & Movie S3). 56% of cells showed no measurable response, maintaining their original speed and showing no or minimal rise (<2%) in Cdc42 activity at their rear despite the stimulation. 21% of cells showed some increase of Cdc42 activity at their rear (>2%) but no engagement of the front which appeared strongly polarized and unyielding, resulting in only cell rear elongation. We termed this class “medium response”. 7% of cells exhibited a “strong response”, transiently flipping their direction of motion, but with Cdc42 polarity and migration reverting back to their original polarization post-stimulation. The remaining 6% of cells reversed. Transient stimulation was significantly less effective in overwriting polarity and reorienting cells as compared to persistent stimulation (6% instead of 47% reversing cells, respectively; Fig. 3b). Notably, the fraction of non-responding cells in the 12-pulse assay was comparable to the fraction of cells that did not reverse upon persistent stimulation (56% and 53%, respectively Fig. 3b). This suggests that cells forming the medium and strong response classes may represent cells with pre-existing conditions that permitted responsiveness and repolarization, but required greater stimulus inputs to completely reverse direction.

**Figure 3:**
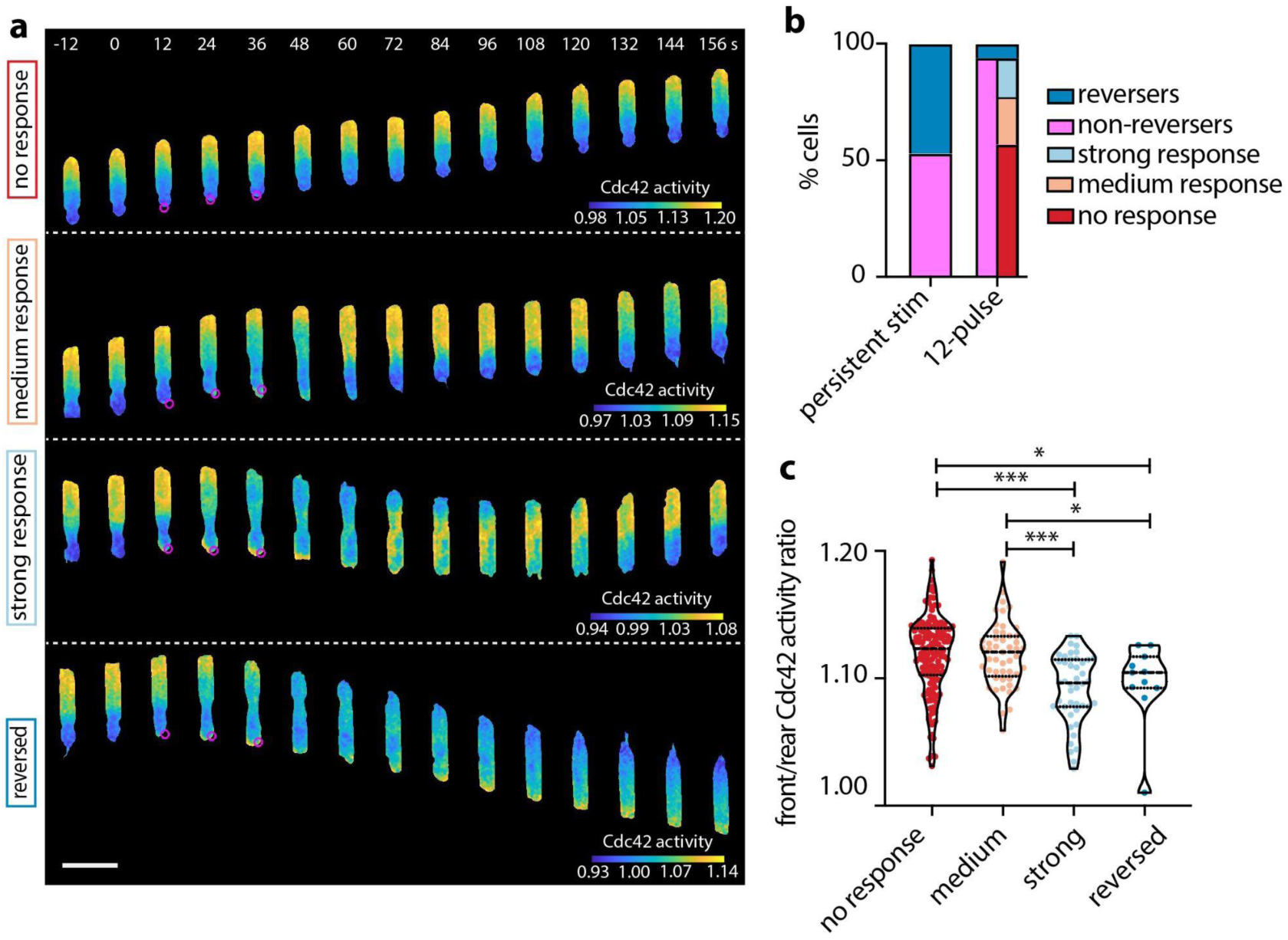
Transient stimulation reveals a variety of distinct cellular responses. **(a)** Representative live-cell imaging snapshots of stimulation experiments with cells expressing parapinopsin and a red/far-red Cdc42 FRET sensor, showing no response (upper panel), medium response, strong response, and a reversed response (lower panel). Cells migrated unperturbed for 60 s prior to starting a transient 12-pulse stimulation at their cell rear (magenta circles). Images captured every 3 s and subsampled for illustration purposes. Scale bar: 25 µm. **(b)** Stacked bar plot of the percentage of cells that exhibited each cellular response with persistent pulsated stimulation (n=336 cells from 36 independent experiments), and with transient 12-pulse stimulation (n=264 cells from 20 independent experiments). **(c)** Violin plot of mean front/rear Cdc42 activity ratio of n=141 non-responders, n=49 medium responders, n=44 strong responders, and n=11 reversers, averaging over 15 s prior to stimulation; p-values of two-sided Wilcoxon rank sum test (*: p<0.05, ***: p<0.001, pairs not shown have p>0.05).

We asked whether we could identify any pre-stimulation variations among cells that exhibited these distinct responses. We found that strong responding and reversing cells were slower migrators with weaker polarization than cells that showed no or medium response (Supplemental Fig. 3a and Fig. 3c). Non-responding cells had the strongest rears (i.e. lowest rear-localized Cdc42 activity) among all classes (Supplemental Figs. 3b-3c). Lastly, front Cdc42 activity was higher in medium compared to strong responding cells (Supplemental Fig. 3d), whose front eventually yielded to the original rear and transiently repolarized.

One possible explanation for why strong responding cells regained their original polarity orientation is that the pre-existing serum gradient might flip back their polarity axis after we switched off the optogenetic stimulus. We therefore examined what would happen if we performed similar experiments for cells moving through channels in a uniform serum environment instead of a gradient, but found no significant differences (Supplemental Fig. 3e), indicating that detection of the pre-existing serum gradient cannot explain why strong responding cells reorient back toward their original direction of motion after optogenetic stimulation ceases.

### A strong cell rear is refractory to receptor inputs

We quantified the average centroid speed traces for the four different classes, namely no response, medium response, strong response and reversed (left to right Fig. 4a), as well as the magnitude and derivative of the Cdc42 activity at each edge (Figs. 4b-4c). The derivative revealed a key difference between the cells in the strong response and the reversed classes. The original rear of strong responding cells showed both a strong positive response to the initiation of stimulation and a prominent negative overshoot post-stimulation (black arrow on the third panel of Fig. 4c). In contrast, reversers showed a lower positive response and no overshoot. Simply put, a cell that overreacted when the stimulation commenced would also overreact when it stopped, emerging as a strong responder. This combination of positive and negative regulation of Cdc42 downstream of receptors was also documented in cell-wide optogenetic stimulation experiments and may reflect temporal processing of input signals^14^.

**Figure 4:**
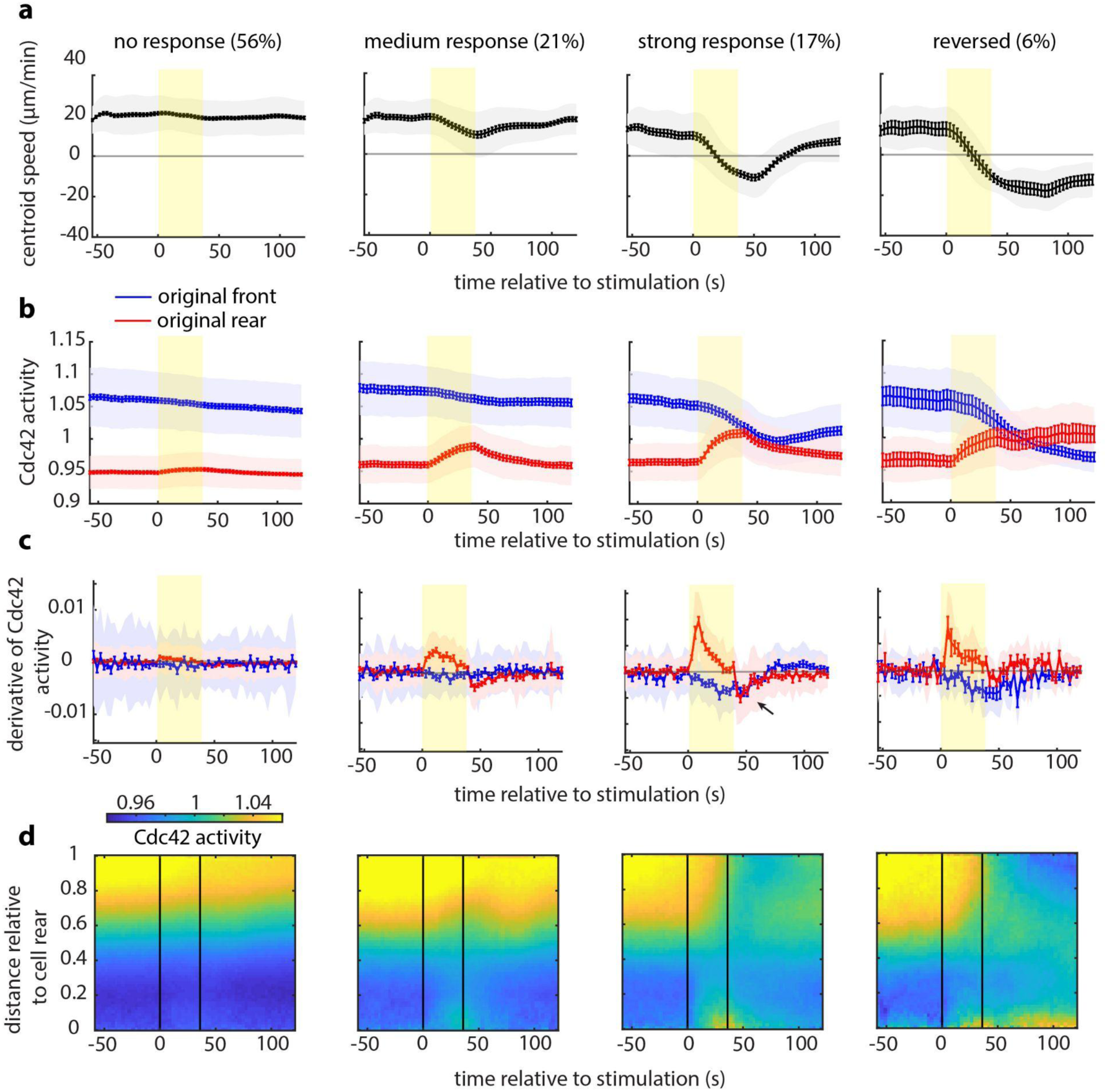
A strong cell rear is refractory to receptor inputs. **(a-c)** Cell centroid speed **(a)**, mean Cdc42 activity **(b)** and derivative of Cdc42 activity **(c)** at the original front (blue) and rear (red) over time for each cellular response (lines: means, shaded regions: SD, error bars: SE). Data are averages from n=141 non-responders, n=49 medium responders, n=44 strong responders, and n=11 reversers from 20 independent experiments. Rectangular yellow shaded region marks the start and end of the 12-pulse stimulation. **(d)** Average kymographic representation of Cdc42 activity as a function of time (x-axis) and vertical position relative to the cell rear (y-axis) for non-responding, medium responding, strong responding and reversing cells. Vertical black lines indicate the start and end of the pulsated stimulation.

Finally, average kymographs for the four distinct cellular responses compactly summarized on a whole-cell level the dynamic spatiotemporal response of Cdc42 activity in each cell class (Fig. 4d). The kymographs further highlighted our most surprising observation: that a strong rear can be refractory to receptor inputs, even at the level of signal transmission to Cdc42. In contrast, there was a clear trend in which a weaker rear can be amenable to receptor input and lead to a measurable whole-cell response. Overall, our observations reveal that the strength of the rear is key in modulating the magnitude of the behavioral response, raising the question of what tunes the rear’s strength in the first place.

### The phosphorylation of myosin regulatory light chain tunes the amenability of the rear to respond to receptor inputs

We sought to explore what molecular pathways might be actively suppressing the rear response to receptor inputs. We considered two main candidates: i) myosin II/RhoA and ii) ERM (ezrin, radixin, and moesin) proteins/cortical actin.

Myosin II and RhoA signaling have been extensively shown to limit membrane protrusion at the cell rear antagonizing front polarity signaling^3, 15, 16^. Myosin motor activity and myosin filament assembly are regulated by the phosphorylation of Ser19 and Thr18 on the myosin regulatory light chain (MRLC)^17–19^. RhoA activates Rho-associated kinase (ROCK) which directly phosphorylates MRLC and inhibits the phosphatase PP1 with its regulatory myosin-binding subunit MYPT1^20^ (Fig. 5a). ROCK inhibition with Y27632 decreases MRLC phosphorylation^21, 22^. Moreover, microtubules have been demonstrated to regulate MRLC phosphorylation by sequestering GEF-H1, a guanine nucleotide exchange factor of RhoA, in various cell types^23–26^. In neutrophils, microtubule destabilization results in activation of RhoA, hyperphosphorylation of MRLC and enhanced myosin contractility^22, 27^. Consequently, if the amenability of the cell rear to respond to receptor inputs is modulated by myosin II/RhoA activity we would expect that Y27632 treatment would reduce the antagonistic effect of myosin II/RhoA and make the rear more amenable to optogenetic reprogramming. Conversely, nocodazole treatment, shown to globally increase RhoA activity and MRLC phosphorylation^22, 28, 29^, would be expected to further enhance the strength of the back polarity module, thus suppressing cell rear response.

**Figure 5:**
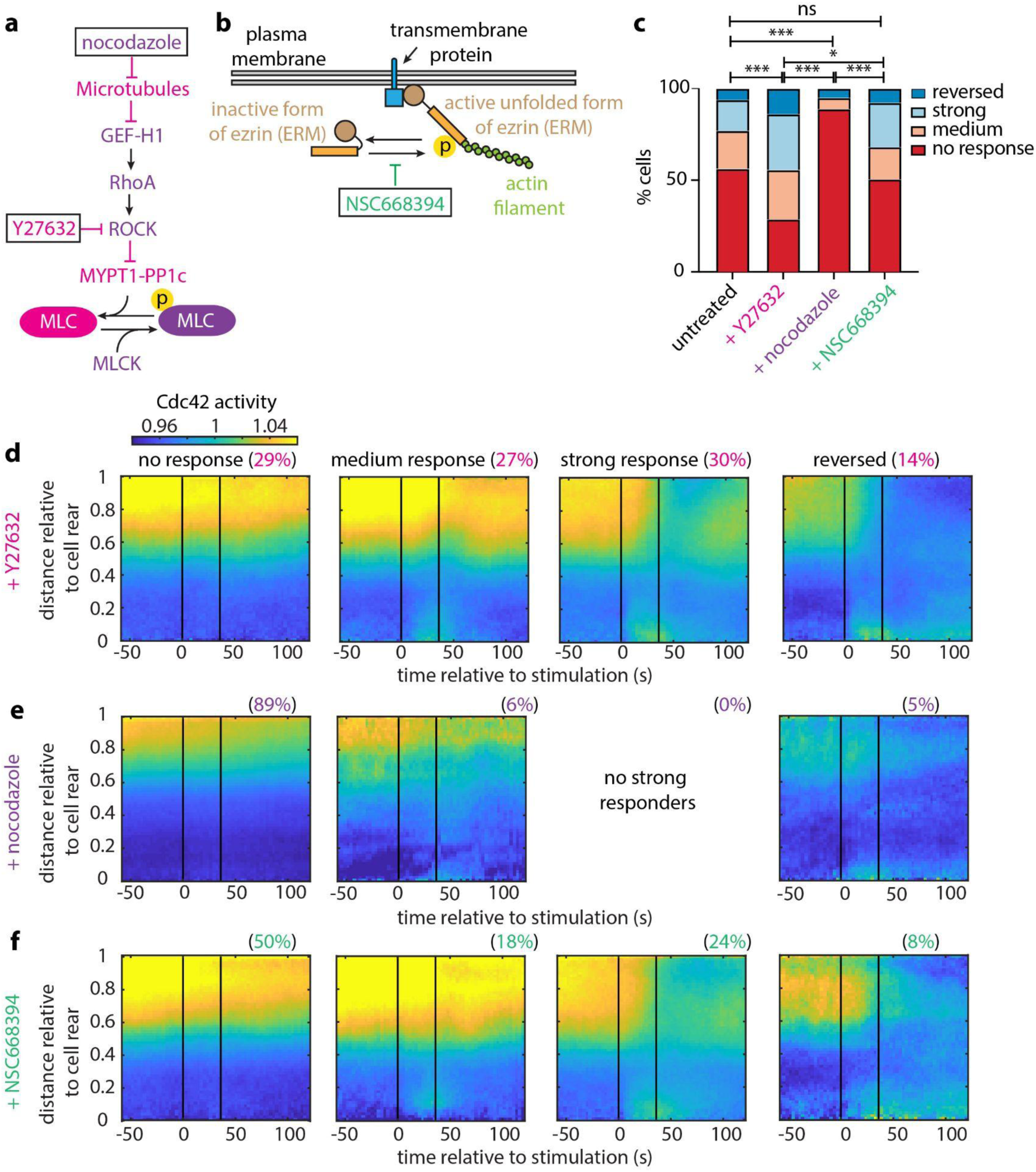
The phosphorylation of myosin regulatory light chain tunes the amenability of the rear to respond to receptor inputs. **(a)** Diagram of the signaling cascade regulating MRLC phosphorylation (adapted from ^22^). A protein or perturbation that increases or decreases MRLC phosphorylation is colored with purple or magenta, respectively. **(b)** Sketch showing how the ERM proteins establish crosslinks between the plasma membrane and cortical actin. **(c)** Stacked bar plots of percentage of cells that showed no response, medium response, strong response and reversed for n=264 untreated cells, for n=115 Y27632-treated cells, n=99 nocodazole-treated cells, and n=91 NSC668394-treated cells (from 20, 7, 7 and 6 independent experiments, respectively); p-values of fisher exact tests (*: p<0.05, ***: p<0.001, ns: p>0.05). **(d-f)** Average kymograph representation of Cdc42 activity as a function of time (x-axis) and vertical position relative to the cell rear (y-axis) for n=107 Y27632-treated cells (d), n=76 nocodazole-treated cells **(e)**, and n=84 NSC668394-treated cells **(f)**. Vertical black lines indicate the start and end of the pulsated stimulation.

ERM proteins have also been implicated in cytoskeletal remodeling and cell migration^30, 31^. ERM proteins transition between an inactive/closed configuration in the cytosol and an active/phosphorylated form that links cortical actin to the plasma membrane^32^ (Fig.5b). Evidence suggests that the ERM protein moesin in its active/phosphorylated form is a key player in neutrophil polarization inhibiting the small GTPases Rac, Rho, and Cdc42 as well as protrusion at the cell rear^33^. Additionally, the persistence of directional migration of neutrophil-like cells towards fMLF gradients has been proposed to depend, at least in part, on the polarization of ERM proteins and moesin towards the cell rear^12^. Specifically, treatment of neutrophils with NSC668394, a quinoline that inhibits ezrin phosphorylation at Thr567 and ezrin-actin binding, has been shown to result in a considerable reduction of directional memory^12^. The above observations made NSC668394 a key perturbation to assess the involvement of ERM proteins in creating a refractory cell rear. Overall, we found that NSC668394 had no significant effect, while Y27632 and nocodozole had highly significant and opposite effects (Fig. 5c), suggesting that the dominant effect on the rear sensitivity is the myosin II/RhoA pathway rather than through ERM proteins.

Specifically, treating cells with Y27632 reduced the number of cells in the non-responding class (Fig. 5c), consistent with what we expected from our hypothesis that myosin II/RhoA could modulate the responsiveness of the rear. Interestingly, the non-responders decreased in number by almost 2-fold as compared to control, the medium responders increased marginally (1.3-fold), and the strong responders and reversers increased by 1.8-fold and 2.3-fold, respectively. The strong and the reversed classes gained more than what would be expected from just depleting the non-responders, suggesting that the Y27632 treatment pushes each cell along the response distribution rather than simply suppressing one particular class. Thus, Y27632 is likely not only making the rear more amenable to optogenetic inputs but also may act on a whole-cell backness component.

In contrast, nocodazole treatment resulted in a striking 1.6-fold increase of the no response class as compared to control (Fig. 5c). Medium responding cells decreased by 3.6-fold, and no cells under this condition exhibited a strong response, whilst the fraction of cells that reversed was comparable that in the control case. Nocodazole’s effect of increasing the fraction of non-responders was exactly what we expected as the converse of the Y27632 finding, further strengthening the evidence of a role for MRLC phosphorylation in tuning the responsiveness of the rear to receptor inputs.

In addition, average kymograph representation of the Cdc42 activity for the tested perturbations showed a consistent trend: non-responding cells were strongly polarized, whereas cells that showed some kind of response were progressively (from medium, to strong, to reversed) more and more weakly polarized (Figs. 5d-5f).

### Myosin II suppresses cellular reorientation and redistributes more slowly than the Cdc42 activity response

Overall, our results pointed to a model in which MRLC phosphorylation regulates the rear response to receptor inputs. In order to visualize myosin dynamics in optogenetic-driven reversals, we generated an HL60 cell line stably expressing fluorescently-tagged MRLC (Myl9-mScarlet) and parapinopsin. We found that the myosin line exhibited a higher percentage of non-responding cells as compared to the Cdc42 line (Fig. 6a). Using flow cytometry, we confirmed that all cells expressed the parapinopsin construct (Supplemental Fig. 4a). This observation led us to hypothesize that the over-expression of MRLC might itself suppress cellular response, consistent with a causative role for myosin II activity in creating a refractory rear.

**Figure 6:**
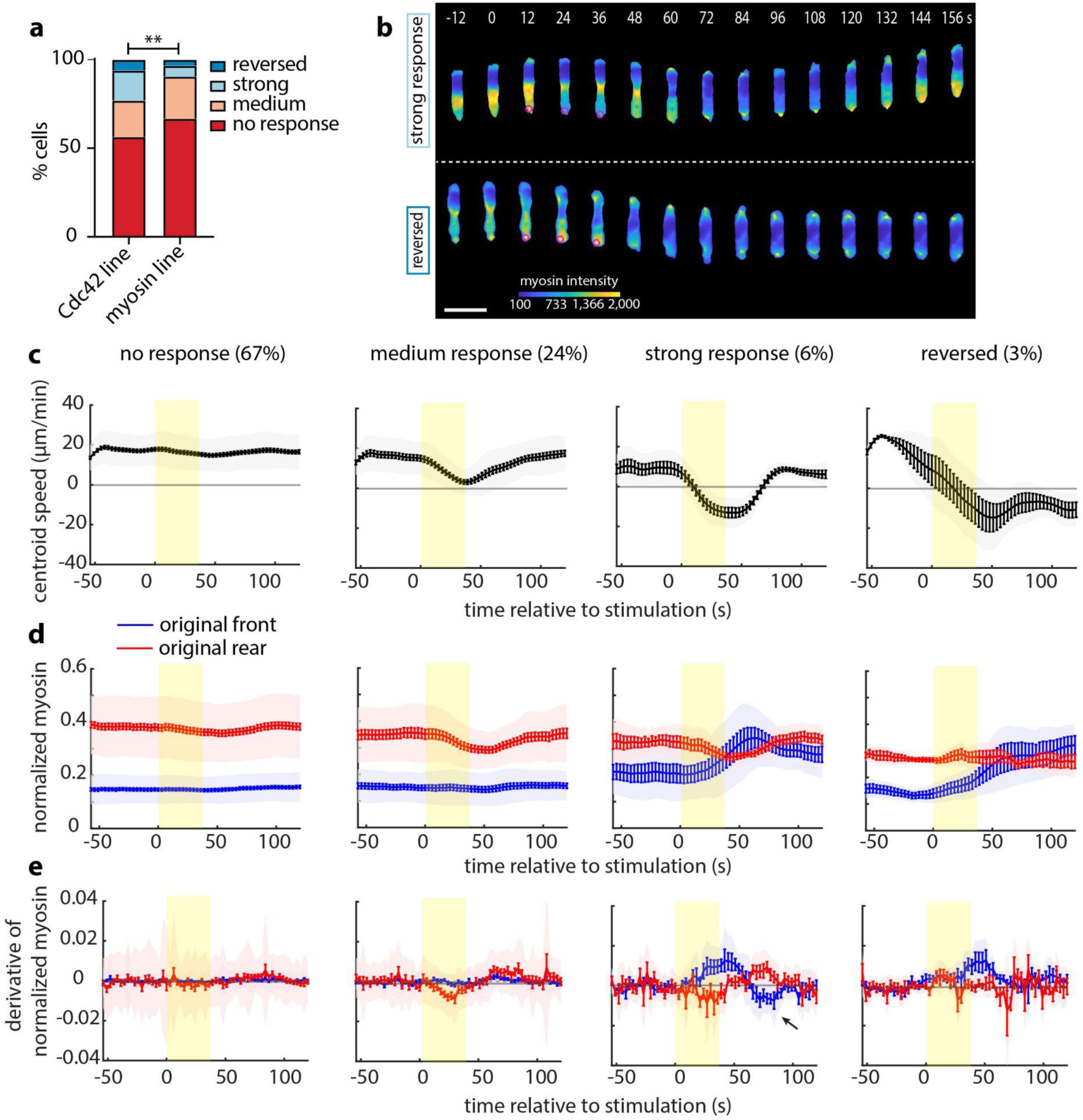
Myosin II suppresses cellular reorientation and lags behind Cdc42 activity response. **(a)** Stacked bar plots of percentage of cells that showed no response, medium response, strong response and reversed for n=264 cells of the Cdc42 line and n=147 cells of the myosin line (from 20 and 8 independent experiments, respectively); p-value of fisher exact test (**: p<0.01). **(b)** Representative live-cell imaging snapshots of stimulation experiments with cells expressing parapinopsin and a myosin light chain sensor/cytosolic tag, showing a strong response (upper panel), and a reversed response (lower panel). Cells migrated unperturbed for 60 s prior to starting a transient 12-pulse stimulation at their cell rear (magenta circles). Myosin intensity pseudo colored to facilitate visualization. Images captured every 3 s, and subsampled for illustration purposes. Scale bar: 25 µm. **(c-e)** Mean cell centroid speed **(c)**, magnitude **(d)** and derivative **(e)** of Cdc42 activity at the original front (blue) and rear (red) over time for each cellular response (lines: means, shaded regions: SD, error bars: SE). Data are averages from n=98 non-responders, n=35 medium responders, n=9 strong responders, and n=5 reversers. Rectangular yellow shaded region represents the start and stop of the pulsated stimulation.

Myosin was polarized to the rear of cells in each response class, but with varying degrees of polarization (Fig. 6b and Supplemental Fig. 4b, also Movie S4). The average centroid speed traces for the four classes (Fig. 6c) were qualitatively identical and temporally close to the ones computed from the Cdc42 line (Fig. 4a), allowing us to draw direct comparisons between Cdc42 activity and myosin dynamics.

We quantified the subcellular normalized myosin II localization for cells in each response class and found that myosin relocalization lags behind Cdc42 activity response (Fig. 6b), suggesting that Cdc42 activity is a faster representation of polarity change. Furthermore, the derivative of the normalized myosin at each edge reveals the same key difference we observed on a Cdc42 activity level between the strong and the reverse response (black arrow on the third panel of Fig. 5e). That is, strong responding cells “overreact” both at the onset and the end of stimulation compared to reversing cells.

Finally, average kymographs of intracellular myosin II further illustrated how myosin over-expression seemed to bias cell behavior, with non-reversing cells having higher myosin levels (Supplemental Fig. 4c). This trend was also captured by comparing the averaged pre-stimulation myosin signal over the entire cell body (Supplemental Fig. 4d). Importantly, the difference was not just in the overall myosin expression level but also in its subcellular distribution, reflecting stronger asymmetry in myosin activity. We quantified the pre-existing subcellular myosin localization (Supplemental Figs. 4e-4g) and found a larger accumulation of myosin at the rear of non-responding cells (Supplemental Fig. 4e). Overexpression may result in increased contractile activity and steeper myosin polarization, rendering the cell rear more refractory to receptor inputs.

### The phosphorylation of myosin regulatory light chain alters the intracellular localization of myosin II and modulates the receptiveness of the rear to respond

Treating the myosin cell line with Y27632 and nocodazole revealed the same trends as noted before with the Cdc42 line: Y27632 significantly reduced the percentage of non-responding cells as compared to control, whereas nocodazole greatly increased it (Fig. 7a). Notably, the proportion of myosin-expressing non-responding cells was higher in all conditions as compared to the Cdc42-expressing cells under the respective treatment conditions, consistent with the hypothesis that the myosin line is less amenable to input simply due to overexpression of the fluorescently-tagged MRLC.

**Figure 7:**
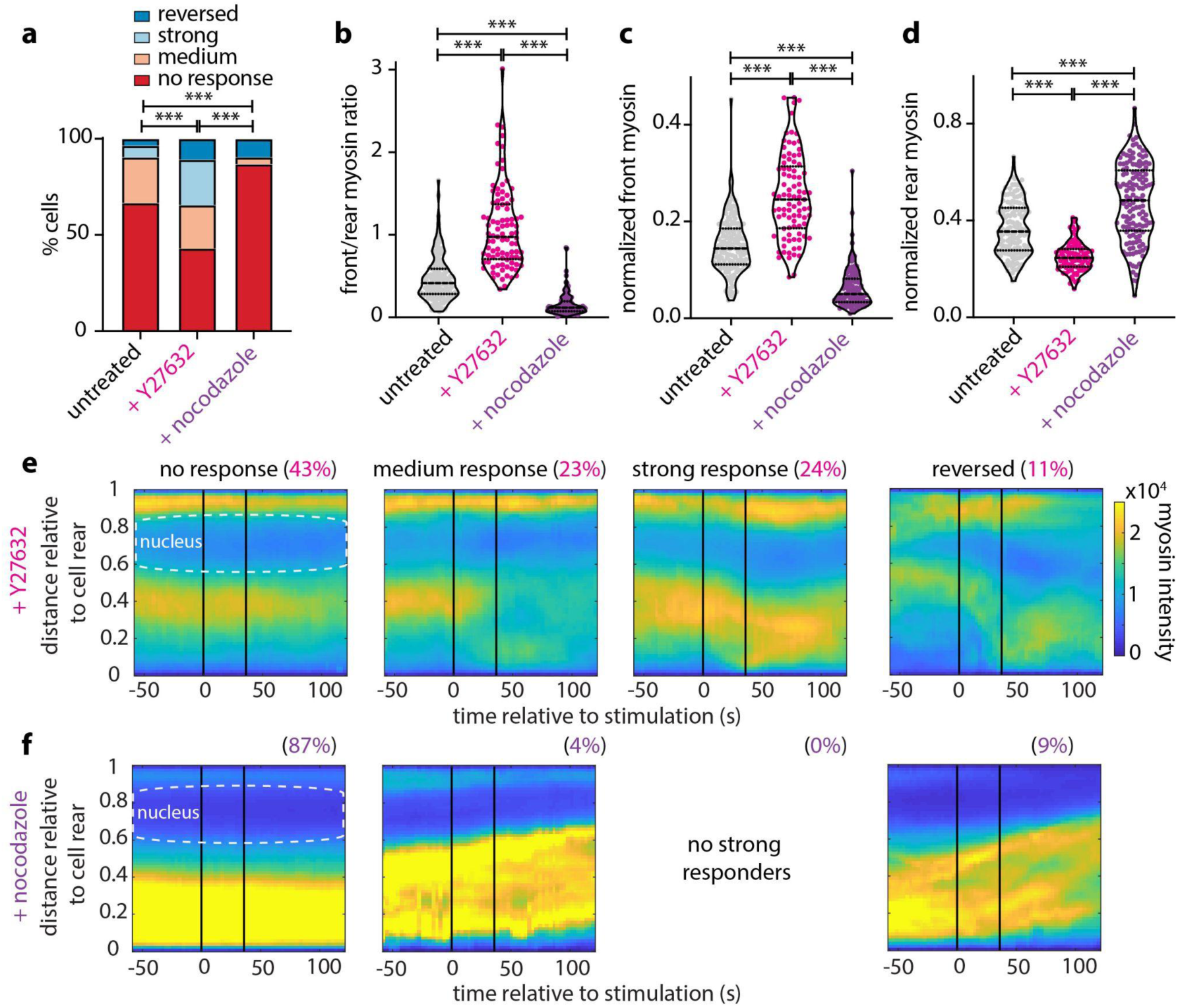
Myosin phosphorylation state alters the intracellular localization of myosin II and tunes the sensitivity of the cell rear. **(a)** Stacked bar plots of percentage of cells that showed no response, medium response, strong response and reversed for n=147 untreated cells, for n=93 Y27632-treated, and n=137 nocodazole-treated cells (from 8, 10 and 8 independent experiments, respectively); p-values of fisher exact tests (***: p<0.001). **(b-d)** Violin plots of mean front/rear myosin ratio **(b)**, normalized front myosin **(c)** and normalized rear myosin **(d)** averaging over 15 s prior to stimulation for the same conditions and cells shown in **(a)**; p-values of two-sided Wilcoxon rank sum test (***: p<0.001). **(e-f)** Average kymograph representation of myosin intensity as a function of time (x-axis) and vertical position relative to the cell rear (y-axis) for cells treated with Y27632 (n=93) **(e)** and nocodazole (n=137) **(f)**. Vertical black lines indicate the start and stop of the pulsated stimulation.

We confirmed that drug perturbations were not affecting myosin expression (Supplemental Fig. 5a). However, the perturbations had measurable effects on subcellular myosin distribution, as quantified by front/rear myosin ratio (Fig. 7b), normalized front (Fig. 7c), rear (Fig. 7d) and middle myosin (Supplemental Fig. 5b), as well as cellular speed (Supplemental Fig. 5c). Y27632 greatly suppressed rear myosin as compared to the control and increased the normalized front myosin, resulting in a more symmetric myosin distribution. Acting opposite, nocodazole dramatically increased rear myosin and suppressed the front myosin pool. Average kymographs of myosin II for Y27632- and nocodazole-treated cells (Fig. 7e-7f, respectively) and kymographs of the difference between each drug treatment and the control confirm the above (Supplemental Fig. 5d-5e). In other words, nocodazole amplifies the myosin distribution asymmetry; whereas Y27632 eliminates it to a great extent. Notably, even upon Y27632 treatment, 43% of tagged MRLC-expressing cells were not responsive, suggesting that ROCK-regulated myosin activity and asymmetry, while being a strong predictive component, is not exclusively regulating the rear responsiveness.

## DISCUSSION

In this study we demonstrated a causative role for myosin II and RhoA activity in creating a stabilized cell rear that is refractory to receptor inputs. We showed that the asymmetric localization of myosin II, being a readout of myosin activity, correlates with the strength of the rear, and that increased myosin II in the cell rear suppresses responsiveness to receptor inputs. Simply put, the higher the myosin levels and the greater the asymmetry, the more likely cells are to be non-responsive. Furthermore, we found that Y27632 and nocodazole perturbations, acting in opposite directions of the RhoA/ROCK/myosin II pathway (through decreasing and increasing the phosphorylation of MRLC, respectively), tune the rear’s ability to respond to new receptor inputs and reorient migrating neutrophils.

Our results extend a longstanding model that antagonism between frontness and backness activities stabilizes polarity and causes asymmetric responses to receptor inputs. However, it had been difficult to distinguish whether this asymmetry was due to feedback-based competition between front and back programs or interference with signal transmission from receptors. Our approach allowed us to surgically examine the latter hypothesis by activating receptors only in the rear of cells without perturbing the cell front in a constrained environment. Rather than simply limiting the ability of new inputs to develop a fully functional front and reverse the cell, we found that a high level of myosin II activity at the cell rear was capable of blocking any detectable response, even at the level of Cdc42 activity, in many cells.

In addition, through transient optogenetic stimulation we discovered a particularly interesting class of cellular response, where cells transiently changed direction of motion and repolarized their Cdc42 activity and myosin but reverted back to their original polarity after stimulation ceased. This strong response is reminiscent of a previously documented cellular behavior in which a subset of neutrophil-like cells retained their directional memory after the removal and later re-introduction of external fMLF chemotactic environment^12^. This directional memory was in part associated with ERM protein moesin, as inhibition through NSC668394 reduced this behavior. In our assay, treatment with NSC668394 did not result in a similar suppression of strong responding cells. This leads us to believe that the strong response in our assay is primarily resulting from the sensitivity to signal onset and removal we previously described.

Finally, by directly measuring Cdc42 activity in optogenetic-driven reversals of cells migrating inside 1-D channels, we discovered that the direction of motion flipped before the polarity of Cdc42 activity flipped. In other words, it is not always the cell edge with more Cdc42 activity that defines the migratory direction. Our analysis may reflect neutrophils also engaging temporal-sensing mechanisms, evaluating how Cdc42 activity dynamically changes at each edge, rather than just comparing absolute or relative magnitudes of activity to dictate cytoskeletal polarization and ultimately migratory direction. A subtle point here is that although Cdc42 is a primary regulator of cell direction and formation of protrusive fronts in neutrophil-like cells^4^, there may be other components that could change faster. Nevertheless, evidence of temporal signal processing has also been shown in neutrophil-like cells when confined to migrate in a 2-D plane under agarose^14^, in primary neutrophils upon fast switching of chemoattractant gradients^34^, and in myeloid cells whose persistent migration to certain intermediate chemokines involves temporal sensing^35^.

Historically, the cytoskeleton has usually been perceived as an effector operating downstream of signaling due to receptor inputs. However, growing evidence highlights that the system can also work the opposite way around, with the cytoskeleton critically influencing receptor activity^36^. Our findings strongly support this more complex view (Fig. 8), revealing a new dimension of the interplay between the cytoskeleton and signal transduction.

**Figure 8:**
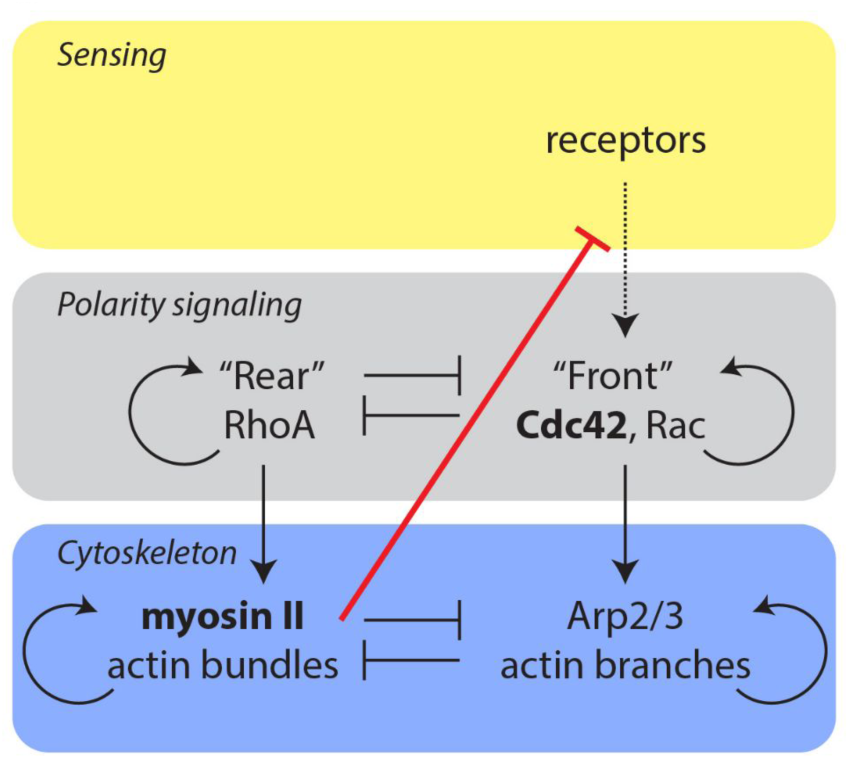
Simplified model of cell polarization. Neutrophil polarization is thought to arise through antagonism between front and rear signaling modules. Information is typically thought to flow from a sensing module containing receptors (yellow layer) to the polarity signaling circuit (grey layer), which guides organization of the cytoskeleton (blue layer). At the front of the cell, Cdc42 and Rac lead to Arp2/3 actin branch formation, while RhoA regulates myosin II actin bundles at the cell rear (vertical arrows). The front and rear signaling modules are mutually exclusive^3^ and are governed by positive feedback loops (semicircular arrows) for self-amplification and polarity maintenance^5–7^. This particular signaling scheme has been demonstrated to be the most robust general motif that gives a stable polarization^37^. Similarly, the cytoskeleton also exhibits positive self-reinforcement and mutual inhibition by the different actin organizations^22, 38–41^. Signaling has usually been perceived to be upstream of the cytoskeleton, informing cytoskeletal responses. Our work revealed that myosin II is capable of blocking any detectable response downstream of receptor activation, even at the level of Cdc42 activity (red line).

## ACKNOWLEDGMENTS

We thank Emel Akdogan for help with the flow cytometry measurements. We thank Tony Y.-C. Tsai and Effie Bastounis for insightful discussions and comments on the manuscript. We thank Bridgette McLaughlin at the UC Davis Cancer Center Flow Cytometry core (supported by P30 CA093373) for technical assistance with cell sorting. This work was supported by the Howard Hughes Medical Institute (J.A.T.), and an NIH Director’s New Innovator Award (DP2 HD094656) to S.R.C. A.H. was a Stanford Bio-X Bowes Fellow, an Onassis Foundation Scholar, and was supported by the Foundation for Education and European Culture (IPEP), the A.G. Leventis Foundation and the Attica Tradition Foundation. Microfabrication work was supported by NIG grant GM092804 to D.I. G.R.R.B. thanks the National Science Foundation Graduate Research Fellowship (Grant Number 1650042) for support.

## AUTHOR CONTRIBUTIONS

A.H., G.R.R.B., S.C. and J.A.T. designed the experiments, A.H. performed the experiments and analyzed the data, A.H., G.R.R.B. and S. C. created software for data analysis and automating experiments, G.R.R.B. and S. C. developed the optogenetic and imaging tools, G.R.R.B. and S.S. created the cell lines, F.E. and D.I. designed the microfluidic devices, A.H., S. C. and J.A.T. interpreted the data, A. H., S. C. and J.A.T. wrote the manuscript. All authors read and approved the manuscript.

## MATERIALS AND METHODS

### Generation of HL60 cell lines stably expressing parapinopsin-Cdc42 FRET sensor and parapinopsin-Myl9

Zebrafish parapinopsina and Cdc42 Tom/Kat FRET sensor plasmids were cloned as previously described^14^. HL60 cell lines were generated by using the 2^nd^ generation lentiviral system to stably insert the parapinopsina gene. Next, the Cdc42 FRET sensor was stably integrated using the piggybac transposon system^42^. The sensor plasmid was co-electroporated at a 1:1 ratio with the piggybac transposase plasmid using the Neon electroporation transfection system (Thermo Fisher Scientific). We used pulse voltage 1350 V and pulse width 35 ms.

The myosin light chain plasmid was generated using multi-part Gibson cloning where iRFP670 fluorescent protein (to enable cytosolic segmentation), a tandem P2AT2A element and mScarlet-I fused to Myl9 were inserted into a lentiviral vector. The plasmid and plasmid maps will be made available on addgene for the published version of this manuscript. The myosin construct was then stably inserted using the lentiviral system. HL60 cells expressing the myosin construct were selected using 10 µg/ml blasticidin (Corning 30-100-RB). The line was sorted for cells highly expressing the myosin light chain (mScarlet-I), and cytoplasmic iRFP670 using a Beckman Coulter Astrios EQ at the UC Davis Flow Cytometry Core Facility.

### Cell culture and differentiation

HL-60 cells were cultured and differentiated into a neutrophil-like state as previously described^43^. Briefly, cells were cultured in RPMI 1640 plus L-glutamine and 25 mM HEPES media (Gibco RPMI 1640 Medium, HEPES, Thermo Fisher Scientific), supplemented with 10% heat-inactivated fetal bovine serum (hiFBS) (Foundation Fetal Bovine Serum, Gemini Bio, heated in a water bath for 40 min at 56 °C to inactivate), 100U/mL penicillin, 10 mg/mL streptomycin, and 0.25 mg/mL Amphotericin B (Gibco Antibiotic-Antimycotic, Thermo Fisher Scientific). Cells were maintained at 37 °C and 5% CO2 in a tissue culture incubator and were passaged every day so as to maintain a cell density close to 5×10^5^ cells/mL. Cell differentiation was achieved by diluting cells in the complete RPMI media at a starting density of 2×10^5^ cells/mL and spiking 1.3% DMSO (Acros 61097). Experiments were performed with cells differentiated for 6 or 7 days.

### Microfluidic device fabrication

Microfluidic devices were prepared as previously described^44^. They consisted of a cell loading channel of 200 µm height and 200 µm width bordered on one side by an array of straight, 944 µm long migratory channels of 6 µm width and 3 µm height. Migration channels were arrayed in groups of three, each triplet connecting to a common 4 nL volume reservoir.

Devices were fabricated using photolithography and soft-lithography approaches. Briefly, photolithography masks were designed using AutoCAD software (Autodesk) and provided in chrome on glass by Front Range Photomask (Lake Havasu City, AZ). The master wafer was prepared by sequentially spin-coating the wafer with two layers of SU-8 negative photoresist (Microchem, Newton, MA) and patterning by exposure to UV light through the two chrome masks. Two cycles were carried out to produce the migration channels at 3 µm height and the loading channels and reservoirs at 200 µm height. To fabricate polydimethylsiloxane (PDMS) devices, the master wafer was used as a replica mold for soft lithography. PDMS (Fisher Scientific, Fair Lawn, NJ) was mixed thoroughly with curing agent at a ratio of 10:1, poured over the master wafer, then degassed for 1 h prior to curing in an oven at 75 °C overnight. Once cooled, PDMS devices were cut from the master wafer and the inlets and outlets punched using a 1 mm biopsy punch (Harris Uni-Core, Ted Pella Inc. Redding, CA). Following treatment with oxygen plasma, devices were then bonded irreversibly to 35 mm glass-bottomed (No. 0 coverslip) gamma-irradiated culture dishes (MatTek Corp. Ashland, MA) by heating to 85 °C for 10 min on a hotplate.

### Microfluidic device priming

To prime the device 20 µL of L-15 media (Gibco) containing 20% hiFBS was pipetted through one of the two loading ports. 10 µL of the same solution was pipetted on top of each loading port. The device was then placed in a vacuum desiccator connected to house vacuum (27 inHg) for 10 min. Upon removal the device was rested for an additional 10 min, until the attractant filled entirely the straight migration channels, connected orthogonally to the central loading channel. The device was then washed twice, by pipetting into a loading port 200 µL of L-15 media. These washing steps removed the attractant from the central loading channel and its passive diffusion from the migration channels into the central loading channel established a serum gradient. To prevent evaporation, 3 mL of L-15 media were added to the glass-bottomed dish to cover the device.

### Retinal preparation

Preparation of retinal stock solutions was performed in a dark room with red-light sources as previously described^14^. In brief, 9-cis-Retinal (Sigma Aldrich R5754) was dissolved in 200 proof ethanol purged (Sigma Aldrich) with argon gas resulting in a concentration of 10mg/mL. Aliquots were stored at −80 °C in amber glass tubes (Sigma Aldrich).

0.1 g of Bovine Serum Albumin (BSA), Fraction V—Low-Endotoxin Grade (Gemini Bio 700-102P) was vigorously mixed in 10 mL of L-15 media (Gibco), to prepare a 1% BSA solution. In darkness, 10uL of retinal stock was diluted to a working concentration of 10 ug/mL by gradually adding the 1% BSA solution (9 x 10 µL, 9 x 100 µL, 2 x 500 µL, 8 x 1 mL) until all 10 mL were used. The final retinal solution was kept in darkness, on a rocker located in a standard cold room, and was left to incubate overnight. Final retinal solutions were used for experiments within 2 days after preparation.

### Retinal incubation and cell loading

3 x 10^5^ differentiated HL60 cells were spun down at 200 g for 5 min and re-suspended in 1 mL of retinal solution to incubate at 37 °C for 1 h. This incubation and all remaining cell handling happened in darkness with red-light sources. Incubated cells were spun down at 200 g for 10 min and were resuspended in ∼20 µL of L-15. Out of this cell suspension, 10 µL were pipetted into the loading port of the device. The device was then transferred on the microscope at 37 °C, where the cells would be imaged. As a control, we performed some experiments without incubating cells with retinal and found that cells were not responsive to stimulation (data not shown).

### Pharmacological perturbations

To explore the role of myosin II/RhoA and ERM proteins/cortical actin in creating a cell back refractory to receptor inputs we pre-treated cells with either 20 µM Υ-26732 (Sigma), or 50 µM Nocodazole (Sigma) or 50 µM NSC668394 (Sigma) for the last 30 min of retinal incubation. Cells were spun down at 200 g for 5 min and were re-suspended in ∼20 µL of L-15 containing the same final drug concentration. Of this cell suspension, 10 µL were loaded in a device that was pre-treated with the same drug concentration. The pharmacological compound was included in both the attractant solution as well as the L-15 solution used for washing, in the same final concentration. For all pharmacological perturbations DMSO was used as a vehicle and 0.15% DMSO As a control, we performed experiments with 0.25% DMSO to match the maximum final DMSO concentration in the drug treatments as controls. We found no significant differences between untreated and DMSO-treated cells (data not shown).

### Fluorescence microscopy

Fluorescence microscopy was performed on a Ti-E inverted Nikon microscope with a XLED1 LED light source for epi-fluorescence illumination. Images were acquired every 3 s on two Andor Zyla 4.2 sCMOS cameras, using a Cairn TwinCam LS image splitter equipped with a dichroic mirror (Chroma ZT594rdc, ∼605 nm edge wavelength) and emission filters to allow simultaneous imaging. The Cdc42 TomKat FRET imaging was performed as previously described^14^. For myosin imaging experiments, we used a 405/488/561/640 nm quad band filter cube (Chroma 91032) that allowed rapid, sequential 407 nm optogenetic stimulation, and simultaneous imaging of mScarlet-I-Myl9 (∼ 561 nm excitation), and cytoplasmic iRFP (∼ 640 nm excitation). Bandpass emission filters were used to eliminate bleedthrough of iRFP into the myosin channel, but some bleedthrough of mScarlet signal into the iRFP channel was unavoidable and tolerated to allow simultaneous imaging. Rapid and precise stage movements were achieved using an ASI stage (MS-2000 Flat-top) equipped with linear encoders. The microscope was controlled through custom-built Matlab software (MathWorks) interfaced with Micro-Manager to automate cell stimulation and time-lapse microscopy protocols, enabling highly reproducible experimental conditions. Cells were imaged with a 60x oil immersion objective (Nikon Apochromat 1.49 NA) and were maintained at 37 °C using a temperature and humidity control unit (OkoLab Microscope Lexan Enclosure). Each experiment was terminated within 2 hours of imaging.

### Stimulation assays

Cell stimulation and imaging was performed using a custom made MATLAB interface for Micro-Manager. Subcellular opsin stimulation was performed using a 407 nm laser (Coherent Cube) with a custom fiber coupling inserted in a FRAP port on the microscope. Both the TomKat dichroic and the dichroic used for myosin imaging can pass 407 nm light, enabling rapid FRAP stimulation without the need to swap cubes. To focus the FRAP module, HL60 cells expressing either the TomKat FRET sensor or the mScarlet-I-Myl9 were loaded in a microfluidic device as previously described. Cells were imaged, as described, to determine the appropriate focal plane. We selected cells, zapped them by setting the laser to 40 ms exposure and 10 % power and imaged them using the FRAP channel. The x-y translation knobs of the FRAP module were adjusted to bring the FRAP spot near the center of the image (around 512 pixel x 512 pixel on a 1024 pixel x 1024 pixel image, where 1 pixel is 0.21 µm). Thereafter, we used the z-adjustment knob to focus the FRAP spot into a tight approximately gaussian-shaped spot. We measured the power of the FRAP laser at the objective using a Thorlabs handheld optical power meter (PM100D) and a microscope slide power sensor (S170C). Cell-stimulation experiments were conducted using ∼1.8 µW power measured at the objective. On each experimental day, pictures of the FRAP spot were collected and averaged to identify the pixel with the maximum FRAP spot intensity. The x-y coordinates of the maximum intensity pixel were saved and later used to dynamically translate the stage, so the maximum activation spot coincided with the desired sub-cellular stimulation location.

Each lane on the microfluidic chip was organized into a grid of x-y coordinates that were imaged consequently. Cells were segmented in real time using a minimum fluorescence threshold and minimum and maximum size thresholds. The target point was computed by determining the cell centroid (in the center stimulation assay) or the cell rear (in all other cases). Determination of the cell rear relied on extracting the cell body boundary from the binary mask and determining the point in the cell perimeter as specified by a target angle (for us 180° in respect to the original orientation of cell movement). For persistent stimulation experiments, UV stimulation and imaging were alternated until the imaging period was over. For the 12-pulse stimulation assays we administered a total of 12 pulses of light, whereas for center stimulation assays we administered a total of 5 pulses. For all assays we maintained the same time interval (3 s) for imaging and stimulation. In all assays cells were allowed to migrate unstimulated for 21 frames and right after imaging the 21^st^ frame, we commenced the sub-cellular stimulation, alternating between imaging and stimulation. This results in turning the receptor off with each image and then rapidly back on again with the following stimulation pulse, as parapinopsina is a G_i_α-family coupled GPCR that is activated by UV light and inactivated by orange light (> 530 nm) (Bell et al. 2021), leading to rapid activation and deactivation cycles. Thus, our stimulation assays are pulsed.

### Camera and illumination corrections

Raw images were corrected for the camera dark-state noise, for differences in the camera chip sensitivity, and for dust in the light path as previously described^14^. Moreover, a gradient in apparent FRET ratio activity was empirically observed from the top to bottom of the TomKat FRET sensor images due to imperfections in the light path. To correct for this gradient, we developed a ratio correction image. Images of unstimulated Cdc42 TomKat FRET sensor HL60 cells loaded in microfluidic channels were collected systematically with different stage positions and the same 60x objective so that at least one cell was imaged on every portion of the camera sensor. We computed FRET ratios using our standard analysis pipeline for each image, and assembled the images into a 3D image stack. We took the median of the FRET ratio over the stack (including only pixels corresponding to cells) to generate a single representative full-field FRET image. To reduce local variability, we then smoothed this image by taking the median over each 24 pixel x 24 pixel block and smoothing using a gaussian filter (sigma=5). We then smoothly resized the result to generate a “ratio correction image” with the same dimensions as the input image. We applied the correction by dividing the FRET ratio images by the ratio correction image. We note that this last correction was not necessary for the Myl9 imaging.

### Cell segmentation and image background subtraction

Raw FRET pair images were registered using the coordinate-mapping strategy described^14^. Cell segmentation was performed on the sum of the aligned FRET donor and acceptor images, to improve signal to noise ratio. First, we conservatively defined background and cell object pixels. A background image was then determined using the median intensity of background pixels in the local neighborhood of each pixel. The background image was subtracted from the sum image. Object edges were enhanced by first smoothing the image using a broader gaussian filter (sigma=5), and then subtracting the smoothed image from the original image. Finally, the cell object binary masks were determined through the Otsu’s threshold method. For each FRET donor and acceptor image we subtracted the background as defined through the above segmentation strategy. Pixels not included in the cell mask were defined as not a number (NAN), so as to eliminate them from downstream analysis. The FRET ratio image was calculated as FRET acceptor divided by FRET donor. We used a similar strategy to register the mScarlet-I-Myl9 and cytoplasmic iRFP pair images for the myosin cell line and to subtract the background.

### Movie processing

For the supplemental movies (Movie S1-Movie S3), each donor and acceptor image was smoothed using a gaussian filter (sigma=2), after subtracting the background as previously described. We note that this smoothing step was only applied to movies and not for any other analysis. The FRET ratio image was calculated once again as FRET acceptor divided by FRET donor (using the smoothed images). The same smoothing strategy was applied for the mScarlet-I-Myl9 signal (Movie S4). The Cdc42 activity ratio for each cell was a bit different and since the relative activity across each cell is most informative, we choose to show cells in Movies S1-S3 with a FRET range from the 1^st^ percentile to the 99^th^ percentile of the corrected FRET ratio images on a movie-by-movie basis. For Movie S4, since differences in myosin expression levels had functional consequences, we displayed myosin expressing cells with fixed bounds from 100 to 2000.

### Data analysis

#### Analysis of the center stimulation assay and threshold assignment for responding cells

We quantified the mean Cdc42 activity over the entire cell body for 60 cells and found a clear increase in Cdc42 activity for most of the cells that started immediately after the initiation of stimulation, peaking at 15 s, right after the last stimulation pulse (Supplemental Fig. 1c). Cdc42 activity returned back to the baseline 15 s later (30 s after the first light pulse). We went on to quantify the relative increase of Cdc42 activity (Supplemental Fig. 1c) by breaking the cellular response into 4 windows (“Control 1”=[-60:-42] s, “Control 2”=[-39:-21] s, “Control 3”=[-18:0] s and a “Peak” window centered at the maximum Cdc42 activity found between 3 and 33 s with a spread ± 9 s around that, to match the window size with the one of the three other windows). For each cell, we quantified the mean value in each of the 4 windows and then computed a “control” response (by dividing the respective means of “Control 2” by “Control 1”), and a “peak response” (by dividing the respective means of “Peak” by “Control 3”). The control distribution is symmetric around the median 0.9993. Leveraging that, we expected half of the inherently non-responding cells in the peak response distribution to have a relative increase less than 0.9993. Using this statistical argument, we estimated that 93% of cells were responsive (close to the 97% of cells that expressed the opsin). Applying a more stringent threshold at 1.008 (which was exceeded in the control ratio by only 5% of cells) provided a lower bound estimate of 70% responding cells. In both estimated thresholds, the percentage of inherently responding cells was significantly higher than the 47% of cells that reversed upon repetitive stimulation, suggesting that the suppressed percentage of reversing cells was due to some other kind of variation among the cells.

#### Analysis of the Cdc42 activity profile

To quantify the pre-stimulus Cdc42 activity profiles (Fig. 1d) we stratified cells as reversers and non-reversers, based on their migratory speed. We assumed and verified that cells are in a steady-state for the 15 s prior to stimulation, so we averaged over that time window to minimize noise in our measurements. Specifically, for each cell we considered the 6 frames prior to stimulation onset, corresponding to 15 s. For each time point we computed a 1-D profile of the Cdc42 activity, centered at the cell centroid and interpolated the Cdc42 activity profile over a fixed number of points (the average cell length) to account for variabilities in cell length. Interpolation was carried out via fitting a cubic smooth spline with the smoothing parameter set at the default value of 0.5 (where 0 fits a straight line and 1 gives a total fit). We computed a mean profile over each cell averaging these 6 interpolated profiles. As a final step, we averaged across all cells in each of the two groups and computed mean profiles (solid lines), standard deviations (shaded regions) and standard error of the means (error bars).

#### Subcellular localization of Cdc42 activity and myosin II

Cell front and rear areas were defined as the 800 points closest to the cell front and rear edge. The 800 points enabled us to capture the entire penetration depth of the Cdc42 activity at the cell front. We note that the order of events (Fig. 2) was qualitatively the same for different penetration depths (800 points, 400 points and 200 points): the stimulated rear responded first, then the front, followed by the cell reversing its direction of migration, before the Cdc42 cross-over point. To define the cell middle, we computed the major axis of the segmented cell body, divided it into three equidistant length segments, and took a rectangular window around the middle part of the cell. For each area of interest (front, rear and middle) we computed the Cdc42 activity as the sum of the donor signal divided by the sum of the acceptor signal in that area. We used the same strategy to identify the front, rear, and middle of Myl9-expressing cells, so that we can directly compare the dynamic re-localization of Cdc42 activity and myosin II (Fig. 6c-6e). To analyze the relative subcellular distribution of myosin, we computed the normalized myosin at the front, rear and middle as the sum of the Myl9 signal in each area of interest divided by the sum of the Myl9 signal in the entire segmented cell body (Figs. 6d-6e, 7e-7f, Supplemental Figs. 4e-g and Supplemental Fig. 5b).

#### Cell speed, length and area analysis

Cell centroid speed was computed by dividing the displacement of the cell’s centroid between two consecutive frames with the frame time interval (3 s). For each cell its speed trace was then smoothed with a local regression using weighted linear least squares and a 2^nd^ degree polynomial model (“loess” or locally weighted smoothing in Matlab). We used a span of 10% of the total number of data points for this smoothing. We then computed the mean cell speed averaging these smoothed speed traces (Figs. 2c, 2g, 4a and 6c).

Cell length was approximated via computing the major axis of the binary cell mask, while cell area was calculated by summing the total number of pixels of the computed binary cell mask. Relative changes in length and area were plotted (Fig. 2d) by dividing the cell length and area for each cell by its respective length and area before stimulation.

#### Determination of the order of events

Temporal determination of the order of events relied on interpolating the time between the closest time-points to key events, namely rear response, front response, cell stalling and Cdc42 activity cross-point. Stimulated rear response was defined as a 1% increase in the Cdc42 activity at the rear, whereas front response was defined as a 1% decrease in Cdc42 activity at the front from the steady-state baseline values for rear and front at t=0 s. Cell stalling was defined as the interpolated time-point where the cell’s centroid speed was zero, and Cdc42 cross-point as the interpolated time-point where the Cdc42 activities of front and rear intersected. This temporal analysis was performed both on average trends (Figs. 2a-2c) as well as on a single-cell level (Figs. 2e-2f).

#### Stratification into cellular responses

Cells were stratified into reversers and non-reversers based on whether they responded to the persistent optogenetic stimulation by reversing their direction of migration and Cdc42 activity axis (Fig. 1c-1g). Transient stimulation resulted in additional cellular responses, namely no response, medium response, strong response, and stably reversing cells (Figs. 3-5). We classified “no response” cells as cells that showed no or minimal rise (<2%) in Cdc42 activity at their rear, when compared to the pre-stimulation rear activity (averaging over 15 s prior to the initiation of stimulation). Medium responding cells were cells that showed some increase of Cdc42 activity at their rear (>2%) but no engagement of the front (difference between minimum front Cdc42 activity and maximum back Cdc42 activity > 0.03), often resulting in cell body elongation. Strong responding cells were classified as cells that showed engagement of both the rear and the front, but not stably reversing (many of these cells transiently reversed direction but flipped back to their original direction after the stimulus ended). Reversed cells were defined as previously described. To extend this analysis to the myosin cell line, the method had to be adjusted, given the lack of a Cdc42 signal reporter in that line (Figs. 6-7). Similar to before, strong responding and reversed Myl9-expressing cells were classified based on reversing their myosin polarity and direction of migration transiently or stably, respectively (for Y27632-treated cells only migratory speed was considered). Non responding and medium responding Myl9-expressing cells were classified using a combination of migratory speed, morphology and manual curation.

#### Kymograph representation for Cdc42 activity and myosin

For each cell and each time point, a 1-D profile of either Cdc42 activity or myosin was computed from cell front to rear. These 1-D profiles were aligned according to the cell centroid. To account for the fact that the cell length was variable across cells and time points we took these 1-D profiles and aligned them to a fixed average cell length, through interpolation. For each time point we took the interpolated 1-D profiles and computed an average profile from all cells. Interpolation was carried out as described above, via fitting a cubic smooth spline with the smoothing parameter set at the default value of 0.5. Synthesizing the average profiles along time, we constructed kymograph representations (Fig. 4d, Figs. 5d-5f, Figs. 7e-7f, Supplemental Fig. 4c).

### Statistical analysis

For categorical data, day-to-day and experiment-to-experiment variation was consistent with counting noise and the observed standard deviations were reasonably explained by the binomial model in each case. For quantitative signaling, speed, and localization data, cell-to-cell variability was the dominant source of variability. Given the limited and variable number of cells that could be analyzed in each experiment, we felt that individual cells represented the most relevant independent unit for statistical analysis. We were also careful to perform control experiments on each day to avoid potential bias. Based on these considerations, we pooled data from all independent experiments performed. We indicate the number of cells and the number of independent experiments in the Figure Legends. Statistical parameters and significance are reported in the Figures and the Figure Legends. Data are determined to be statistically significant when p < 0.05 by either Wilcoxon rank sum test or Fisher exact test (“ranksum” or “fishertest” in Matlab, respectively).

To depict the % of reversing cells (Fig. 1c and Supplemental Fig. 2d) under persistent stimulation we represented data with bar plots showing the cumulative mean value and an error bar which represents the confidence interval assuming a binomial distribution around the cumulative mean, pulling data from all independent experiments.

For the statistical analysis of the subcellular localization of Cdc42 activity and myosin II as well as the centroid speed, we used violin plots pooling all cells that passed our pre-processing standards of masking and tracking, from all independent experiments (Figures 1e-1f, 3c, 7b-7d and Supplemental Fig. 1a-1c, 3a-3d, 4d-4g, 5a-5c). In each violin plot the dashed line represents the median of the distribution and the dotted lines the 25% and 75% quartiles of the distribution.

To compare among groups (*e.g.* reversing and non-reversing cells or among cellular responses or drug perturbations), we performed non-parametric Wilcoxon rank sum tests.

To depict the % of cells in each response class under titrated stimulation, we used stacked bar plots (Fig. 3b, 5c, 6a, 7a and Supplemental Fig. 3e). We performed the Fisher exact test on the distributions as a whole to examine whether at least one of the classes differed between compared pairs (p-values are reported on the Figures). In addition, we computed the Fisher exact for each cellular response class between conditions (*e.g.* comparing the reversers in untreated cells vs Y27632-treated cells etc.) to probe which classes significantly changed. Based on those two considerations, we highlighted the significant difference in the main text.

### Plotting

Plotting was performed using MATLAB R2018a (MATHWORKS) and GraphPad Prism 8.

---

**Supplemental Figure 1:**
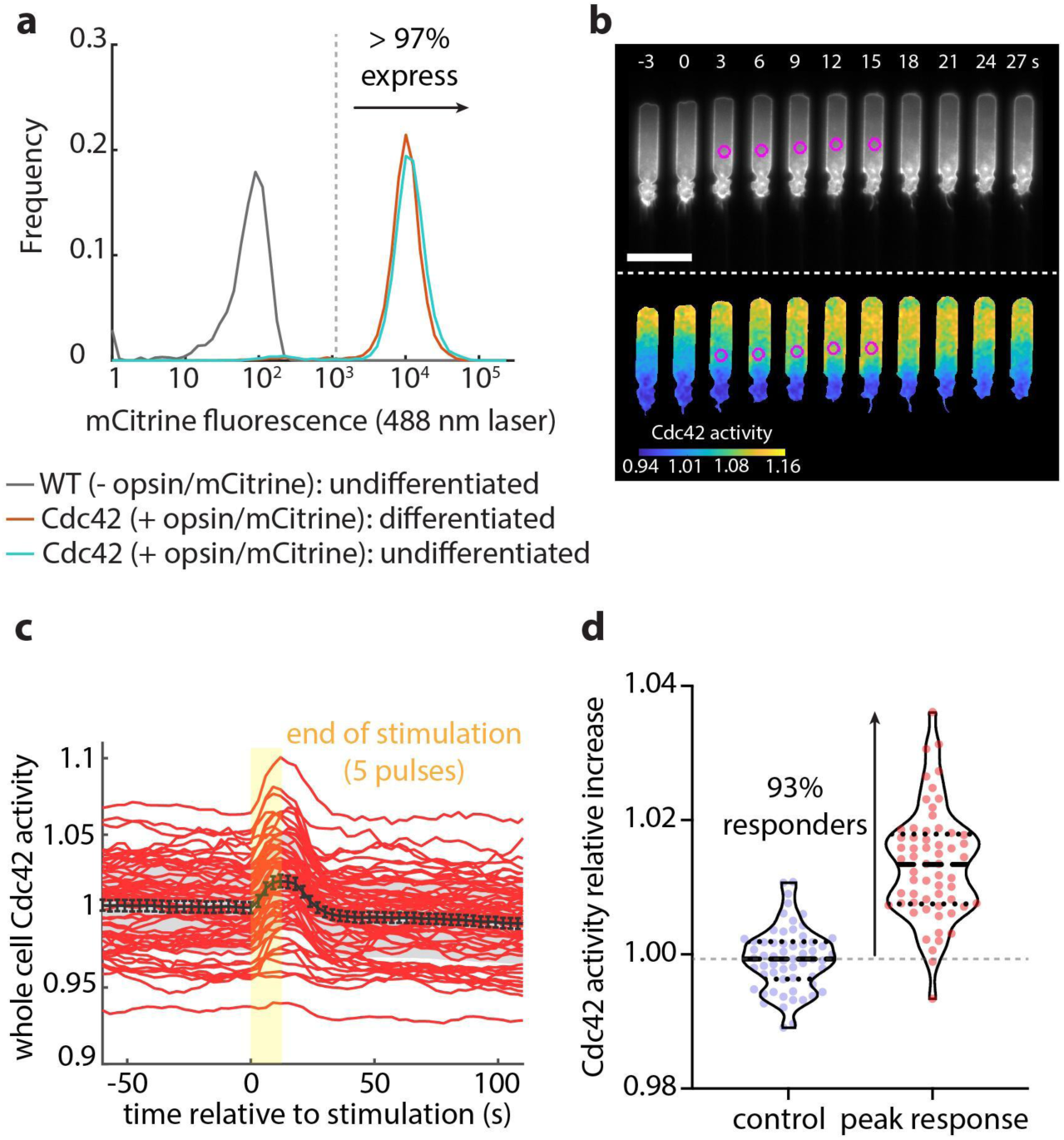
Flow cytometry and center stimulation assay reveal that almost all cells are expressing the opsin receptor and that activated receptors signal to Cdc42. **(a)** Flow cytometry measurement for mCitrine fluorescence in differentiated (red) and undifferentiated (cyan) HL60 cells expressing parapinopsin (tagged with mCitrine) and a red/far-red Cdc42 FRET sensor. Wild type cells (grey) not expressing the opsin used as control. **(b)** Live-cell imaging snapshots of a representative center stimulation experiment on an HL60 cell expressing parapinopsin and the Cdc42 FRET sensor. Cells migrate unperturbed for 60 s before administering 5 pulses at their centroid (magenta circles). Upper and lower panels show the registered sensors (grey scale) and the computed Cdc42 activity, respectively. Images captured every 3 s and subsampled for illustration purposes. Scale bar: 25 µm. **(c)** Mean Cdc42 activity averaged over the entire cell body over time (n=60 cells from 6 independent experiments) stimulated 5 times at their centroid (red lines: individual cells, black line: mean, grey shaded region: SD, error bars: SE). Rectangular yellow shaded region represents the start and end of the 5-pulse stimulation. **(d)** Violin plots of the relative increase of Cdc42 activity over a control zone and over the peak response zone for the same cells shown in **(c)**. Dashed grey line represents the threshold as defined by the median of the control response distribution.

**Supplemental Figure 2:**
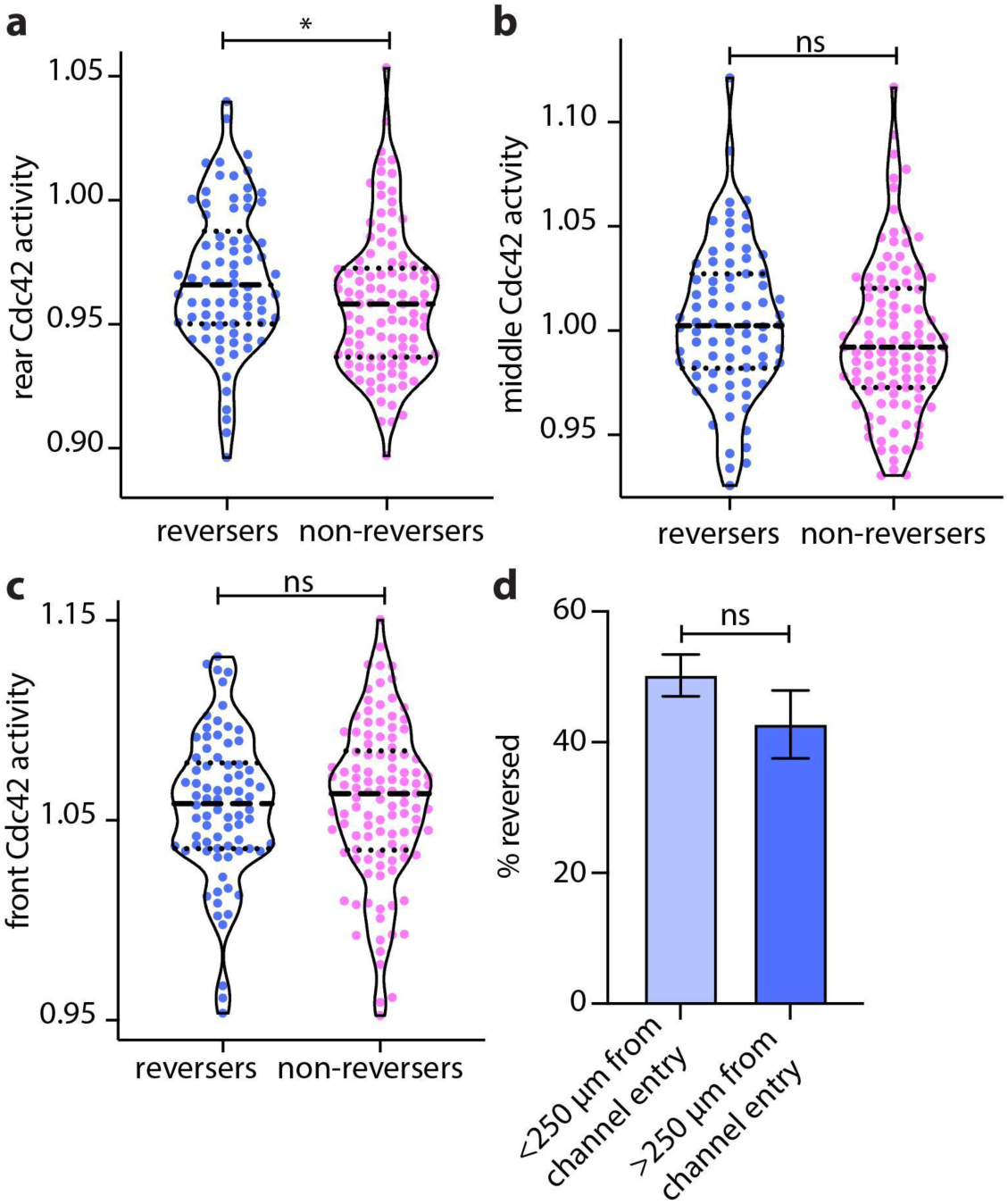
Subcellular analysis reveals that non-reversers have a stronger rear as compared to reversing cells. **(a-c)** Violin plots of mean cell rear **(a)**, cell middle **(b)**, and cell front **(c)** Cdc42 activity of n=78 reversers and n=110 non-reversers, averaging over 15 s prior to initiating persistent rear stimulation; p-values of two-sided Wilcoxon rank sum test (* represents p<0.05, ns represents p>0.05). **(d)** Bar plot of the percentage of cells that reversed stratified by the distance from the channel entrance; closer to the channel entry: distance from entry <250 µm (n=247 cells), farther from channel entry: distance>250 µm (n=89 cells). Error bars represent confidence intervals assuming a binomial distribution around the cumulative mean of each group.

**Supplemental Figure 3:**
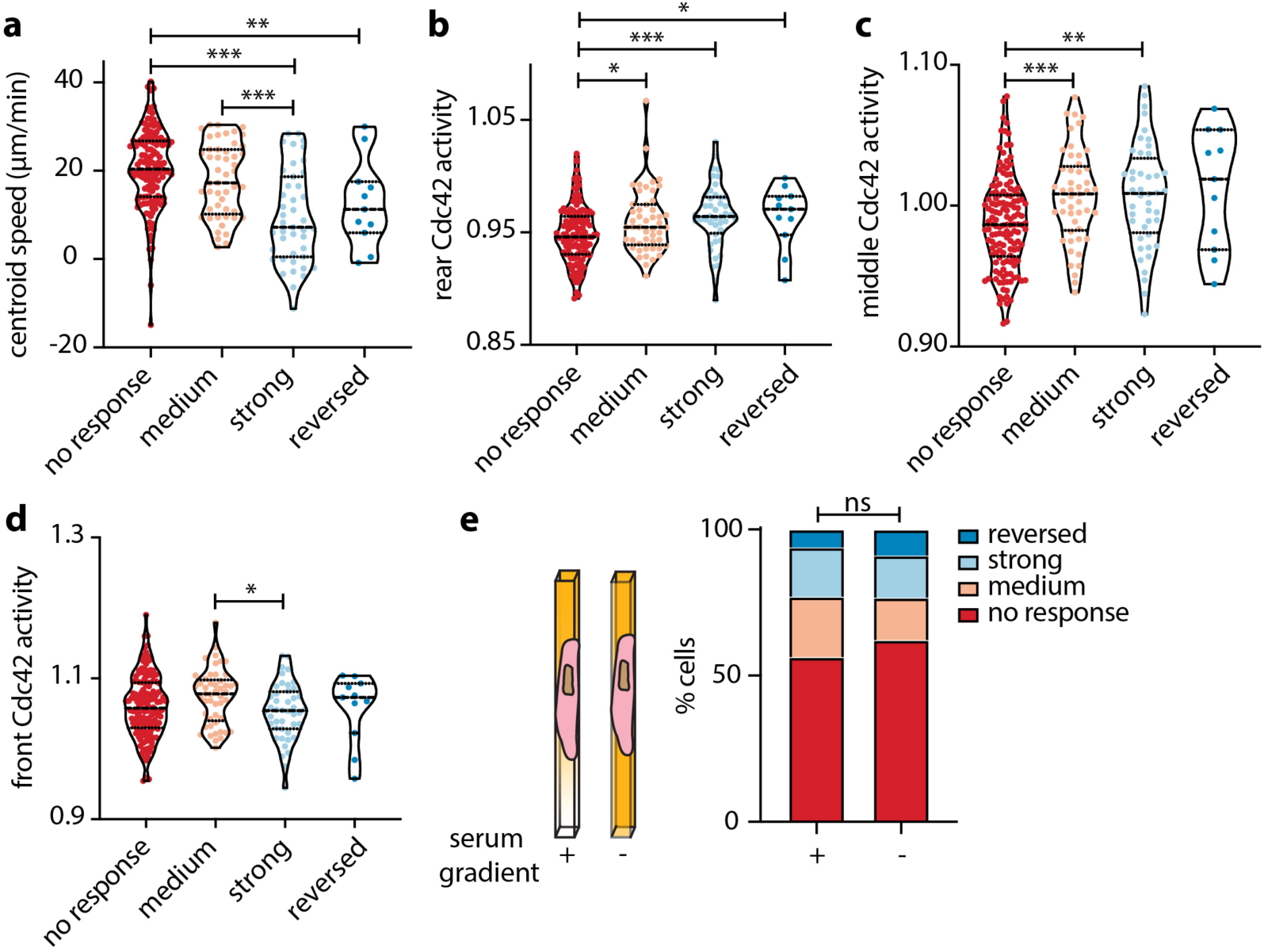
Cdc42 activity quantification reveals that behavioral responses are, in part, due to pre-existing variation. **(a-d)** Violin plots of mean centroid speed **(a)**, and mean cell rear **(b)**, cell middle **(c)**, and cell front **(d)** Cdc42 activity of n=141 non-responders, n=49 medium responders, n=44 strong responders, and n=11 reversers, averaging over 15 s prior to initiating 12-pulse stimulation; p-values of two-sided Wilcoxon rank sum test (*: p<0.05, **: p<0.01, ***: p<0.001, pairs not shown have p:>0.05). **(e)** Stacked bar plots of the percentage of cells that showed no response, medium response, strong response and reversed for n=264 cells that migrated up a serum gradient and for n=101 cells that migrated in a homogeneous serum environment (from 20 and 6 independent experiments, respectively), fisher exact test revealed no significant difference between the two conditions (p>0.05).

**Supplemental Figure 4:**
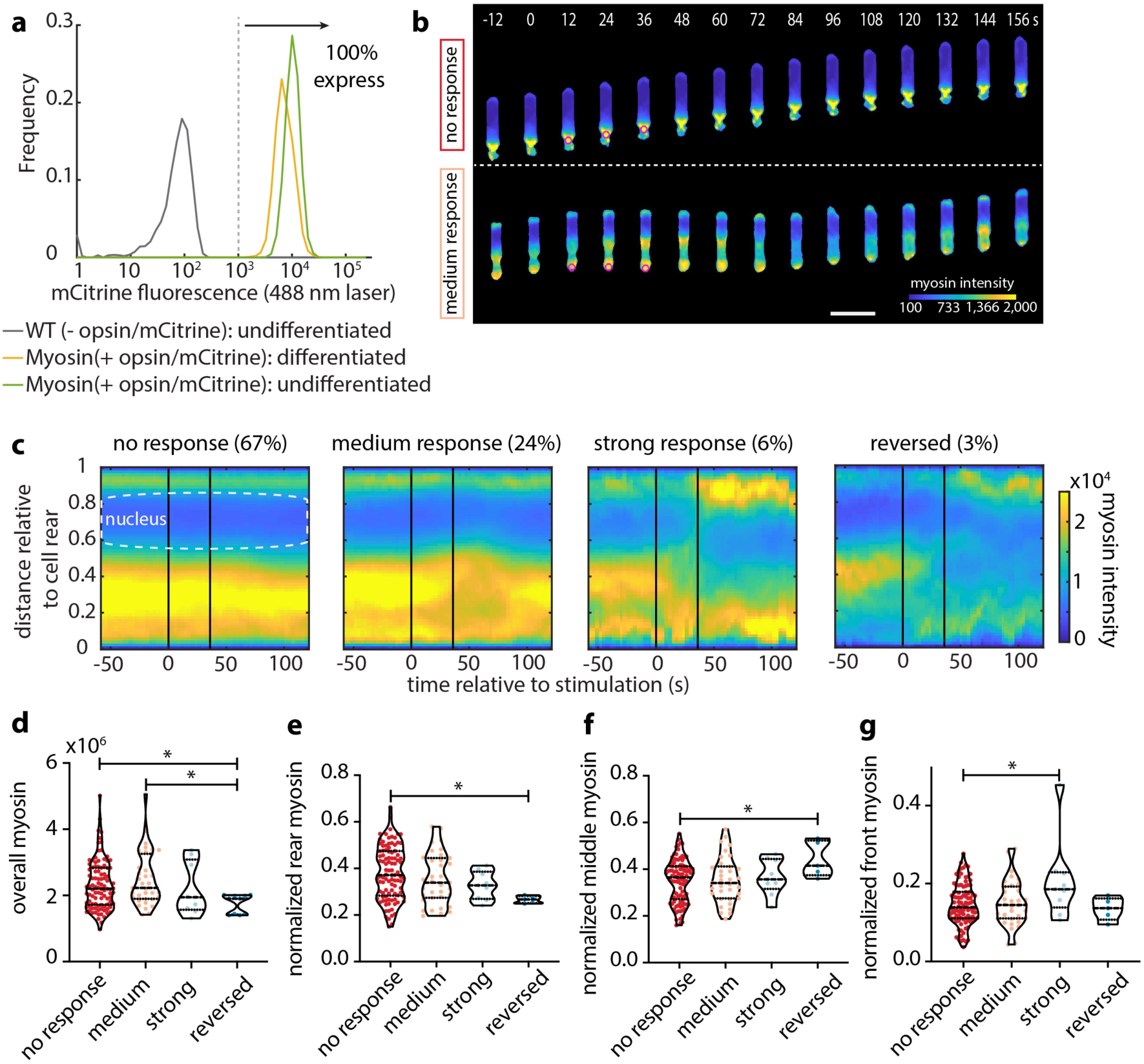
Myosin quantification supports the idea that cellular responses are resulting from pre-existing variation. **(a)** Flow cytometry measurement for mCitrine fluorescence in differentiated (red) and undifferentiated (purple) HL60 cells expressing parapinopsin (tagged with mCitrine), Myl9 and a cytosolic tag. Wild type cells (grey) not expressing the opsin used as control. **(b)** Live-cell imaging snapshots of stimulation experiments with cells expressing parapinopsin and a myosin light chain sensor/cytosolic tag showing no response (upper panel) and a medium response (lower panel). Cells migrated unperturbed for 60 s prior to starting a transient 12-pulse stimulation at their cell rear (magenta circles). Myosin intensity pseudo colored to facilitate visualization. Images captured every 3 s and subsampled for illustration purposes. Scale bar: 25 µm. **(c)** Average kymograph representation of myosin intensity as a function of time (x-axis) and vertical position relative to the cell rear (y-axis) of n=147 cells from 8 independent experiments stratified as non-responding, medium responding, strong responding and reversing cells. Vertical black lines indicate the start and end of the pulsated stimulation. **(d-g)** Violin plots of mean overall myosin **(d)**, mean normalized rear myosin **(e)**, mean normalized middle myosin **(f)**, and mean normalized front myosin **(g)** of n=147 cells, averaging over 15 s prior to initiating 12-pulse stimulation; p-values of two-sided Wilcoxon rank sum test (*: p<0.05, pairs not shown have p>0.05).

**Supplemental Figure 5:**
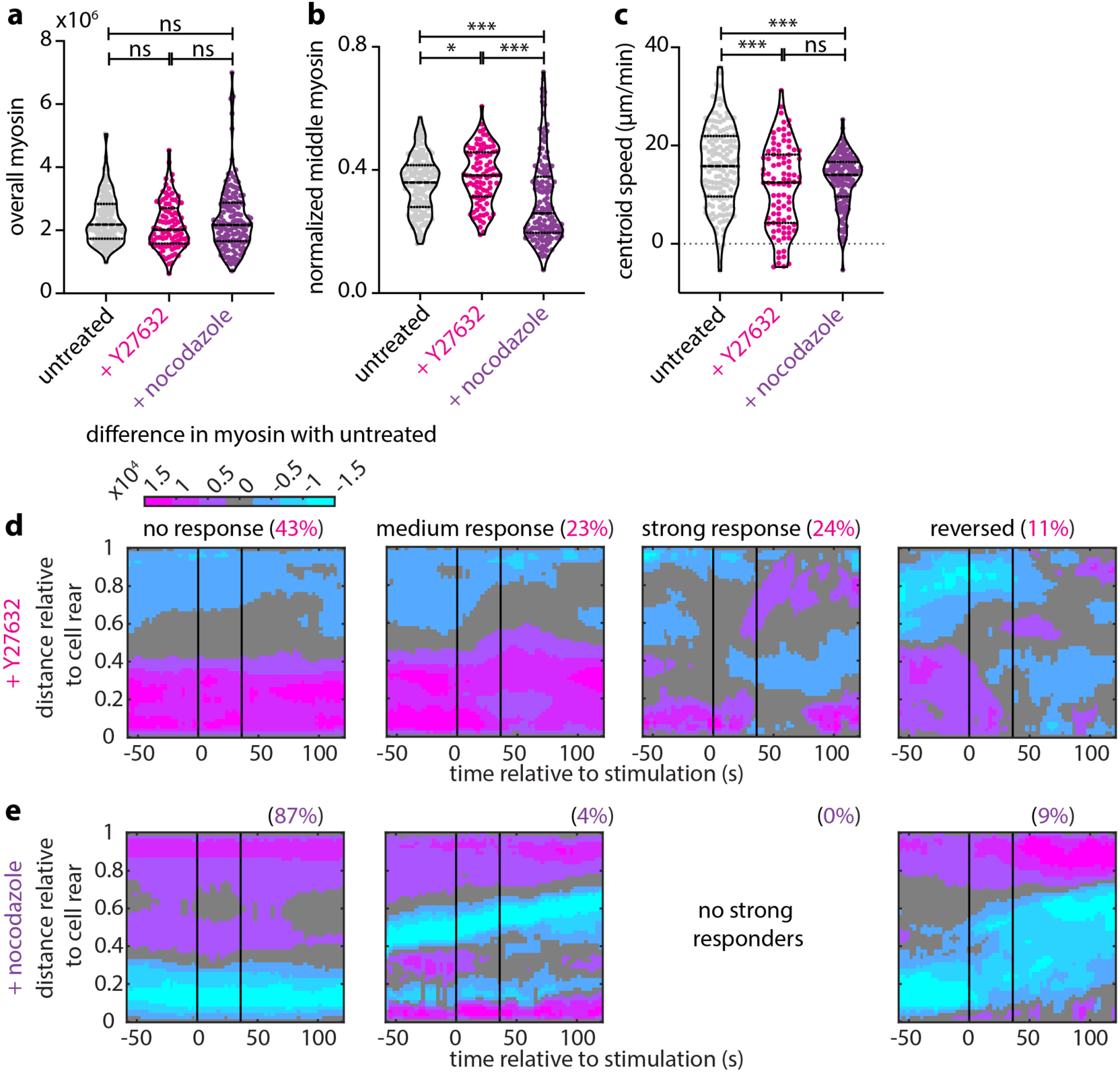
Intracellular myosin localization depends on phosphorylation of myosin regulatory light chain. **(a-c)** Violin plots of mean overall myosin **(a)**, mean normalized cell middle myosin **(b)**, and mean centroid speed **(c)** for n=147 untreated cells, for n=93 Y27632-treated, and n=137 nocodazole-treated cells (from 10 and 8 independent experiments, respectively), averaging over 15 s prior to initiating 12-pulse stimulation; p-values of two-sided Wilcoxon rank sum test (*: p<0.05, ***: p<0.001, ns: p>0.05). **(d-e)** Average kymograph representation of the difference in myosin intensity as a function of time (x-axis) and vertical position relative to the cell rear (y-axis) between n=147 untreated cells and n=93 Y27832-treated **(d)**, and n=137 nocodazole-treated cells **(e)**, stratified as no responders, medium responders, strong responders and reverses (left to right). Vertical black lines indicate the start and end of the pulsated stimulation.

**Movie S1: A cell reversing its direction of motion under persistent optogenetic stimulation at the cell rear.** A successful reversal of an HL60 cell expressing parapinopsin and the Cdc42 FRET sensor. The cell migrated unperturbed for 60 s before administering persistent optogenetic stimulation at its rear (magenta circle). On the left we show the registered sensors (grey scale) and on the right the computed Cdc42 activity. Images captured every 3 s. Scale bar: 25 µm.

**Movie S2: A cell responding on the level of Cdc42 when stimulated at its center.** Representative center stimulation experiment on an HL60 cell expressing parapinopsin and the Cdc42 FRET sensor. The cell migrated unperturbed for 60 s before administering 5 pulses at their centroid (magenta circle). On the left we show the registered sensors (grey scale) and on the right the computed Cdc42 activity. Images captured every 3 s. Scale bar: 25 µm.

**Movie S3: Transient stimulation at the cell rear results in four distinct cellular responses.** Transient stimulation experiment on four different cells expressing parapinopsin and the Cdc42 FRET sensor showing no response, medium response, strong response and reversed (left to right). Cells migrated unperturbed for 60 s prior to starting a transient 12-pulse stimulation at their cell rear (magenta circle). On the left we show the registered sensors (grey scale) and on the right the computed Cdc42 activity. Images captured every 3 s. Scale bar: 25 µm.

**Movie S4: Myosin sub-behaviors under transient stimulation at the cell rear.** Transient stimulation experiment on four different cells expressing parapinopsin and a myosin light chain sensor/cytosolic tag showing no response, medium response, strong response and reversed (left to right). Cells migrated unperturbed for 60 s prior to starting a transient 12-pulse stimulation at their cell rear (magenta circle). Raw myosin intensity on the left and pseudo colored on the right to facilitate visualization. Images captured every 3 s. Scale bar: 25 µm.

